# Programmable mammalian translational modulators by CRISPR-associated proteins

**DOI:** 10.1101/2021.09.17.460758

**Authors:** Shunsuke Kawasaki, Hiroki Ono, Moe Hirosawa, Takeru Kuwabara, Hirohide Saito

**Author notes:** Co-first authors, equally contributed (Shunsuke Kawasaki, Hiroki Ono, Moe Hirosawa).

## Abstract

The complexity of synthetic genetic circuits relies on repertories of biological circuitry with high orthogonality. Although post-transcriptional circuitry relying on RNA-binding proteins (RBPs) qualifies as a repertory, the limited pool of regulatory devices hinders network modularity and scalability. Here we propose CaRTRIDGE (Cas-Responsive Translational Regulation Integratable into Diverse Genomic Engineering) to repurpose CRISPR-associated (Cas) proteins as translational modulators. We demonstrate that a set of Cas proteins are able to repress (OFF) or activate (ON) the translation of mRNAs that contain a Cas-binding RNA motif in the 5’-UTR. We designed 81 different types of translation OFF and ON switches and verified their functional characteristics. Many of them functioned as efficient translational regulators and showed orthogonality in mammalian cells. By interconnecting these switches, we designed and built artificial circuits, including 60 translational AND gates. Moreover, we show that various CRISPR-related technologies, including anti-CRISPR and split-Cas9 platforms, can be repurposed to control translation. Our Cas-mediated translational regulation is compatible with transcriptional regulation by Cas proteins and increases the complexity of synthetic circuits with fewer elements. CaRTRIDGE builds protein-responsive mRNA switches more than ever and leads to the development of both Cas-mediated genome editing and translational regulation technologies.

## Introduction

In all living organisms, highly integrated molecular machinery regulates gene expressions at each step of the central dogma. Fine-tuned control of gene expressions at multiple layers (e.g., transcription, translation, and post-translation) diversifies cellular identity and coordinates multi-cell systems during development, while at the same time its disruption leads to disease^1,2^. Imitating such control holds great potential to engineer cells for biotechnology and medical applications.

In recent years, synthetic biology has made momentous advances in programming cellular behaviours by exploiting DNA, RNA, and proteins in artificial gene regulatory networks^3^. Synthetic biologists have redesigned natural genetic switches to make programmable genetic devices and circuits that evoke silicon-based electronic devices, including band-pass filters^4^, Boolean logic gates^5–7^, arithmetic circuits^5^, oscillators^8^, and memories^9,10^. Transcriptional regulators, such as zinc finger protein fused with transcription factors (ZF-TFs)^11^, transcription activator-like effector (TALE)^12^ proteins, Clustered Regularly Interspaced Short Palindromic Repeats and CRISPR-associated (CRISPR-Cas) systems^6^, recombinases^13^, and meganucleases^14^, are the most well-studied and characterized bio-modules and have been tested for their composability in artificial networks. Recent studies have also succeeded at exploiting post-translational modules, like proteases and protein-protein interactions, as genetic switches^15,16^.

Translational regulators with RNA-binding proteins (RBPs) are one of the most promising and well-exploited synthetic genetic switches for post-transcriptional circuits^17–22^. This approach is DNA-independent, which provides safer cell programming means due to the low risk of genomic damage^18,19^. Additionally, RBP-based translational regulators are DNA-encodable and have been coupled with transcriptional regulators to build designer cells with high-computational potential^5^. However, due to the limited number of available RBPs capable of regulating translation, translation control modules cannot be well standardized, which prevents the implementation of post-transcriptional circuits into mammalian cells.

We hypothesized that CRISPR-associated (Cas) proteins^23,24^ are good candidates to generate RBP-mediated translational controllers for the following reasons. (1) Cas proteins are guided by their corresponding crRNAs (CRISPR RNA) or sgRNAs (single guide RNA: chimeric crRNA and trans-activating crRNA (tracrRNA)). Therefore, Cas proteins can be repurposed as translational controllers by binding to the crRNA or sgRNA embedded in the untranslated regions (UTRs) of messenger RNAs (mRNAs)^25^. (2) Many Cas proteins have been shown to bind to corresponding RNAs in mammalian cells with high specificity and low cytotoxicity, which was confirmed by their catalytic activities. (3) The CRISPR-Cas system is a prokaryotic adaptive immune system^26^. Therefore, its RNA-protein (RNP) interactions may be orthogonal to endogenous RNP interactions in mammalian cells. (4) Due to huge efforts in the CRISPR field, new Cas proteins are being discovered regularly, which will increase the number of potential translational regulators rapidly.

Here we show a new methodology, CaRTRIDGE (<u>Ca</u>s-<u>R</u>esponsive <u>T</u>ranslational <u>R</u>egulation Integratable into <u>D</u>iverse <u>G</u>enomic <u>E</u>ngineering), that repurposes various Cas proteins as translational repressors and activators in mammalian cells. We found that a variety of Cas proteins post-transcriptionally repressed the expression of target mRNAs with high efficiency (OFF switch). They also functioned as translational activators (ON switch) if using a translation-inverter module that converts translational repressors to activators^27^. Moreover, these Cas proteins showed high orthogonality to regulate the translation of target mRNAs, making it possible to construct synthetic translational logic circuits including 60 different AND gates. Cas-mediated translational switches can be combined with conventional transcriptional regulation and reduce the number of elements for the construction of synthetic arithmetic circuits. We also demonstrate that existing techniques for CRISPR-based genome engineering, including conditional transcriptional regulation using anti-CRISPR and split-Cas9, can be repurposed for translational regulation, providing multi-layered gene regulatory systems using Cas family proteins.

## Results

### Repurposing CRISPR-associated proteins as translational modulators

We first tested *Streptococcus pyogenes* Cas9 (SpCas9), which is a well-characterized Cas9 that has been widely used for genome editing^28–30^. To investigate the translational repression ability of SpCas9, we designed SpCas9-responsive mRNA encoding EGFP (SpCas9-responsive EGFP-OFF switch) by inserting sgRNA for SpCas9 (referred to as Sp_gRNA) in the 5’-UTR (Figure 1A, left). We co-transfected a switch-expressing plasmid (switch plasmid) and a SpCas9-expressing plasmid (trigger plasmid) into HEK293FT cells along with an iRFP670-expressing plasmid as a reference (reference plasmid) (Supplementary Figure 1). In principle, SpCas9 should bind to Sp_gRNA in the mRNA and repress translation (Figure 1A, right). We observed that EGFP expression from the switch plasmid with Sp_gRNA in the 5’-UTR was efficiently repressed by SpCas9 induction by flow cytometry (more than 80% repression, Figure 1B and Supplementary Figure 2). To eliminate the possibility that cleavage of the switch plasmid by the DNA-cleaving activity of SpCas9 reduced the reporter EGFP expression, we tested several SpCas9 mutants for their repression activity: SpCas9 nickase (D10A), nuclease-null dCas9 (D10A and H840A), and Cas9 without a nuclear localization signal (ΔNLS)^28,29,31^. These mutants repressed the expression of EGFP with a similar efficiency as wild-type SpCas9 (Supplementary Figure 3A), indicating that the reporter repression was not caused by the DNA cleaving activity of SpCas9. We also examined the effect of AcrIIA4, an anti-CRISPR (Acr) protein that inhibits DNA-binding by SpCas9, on the reporter expression^32^. The co-transfection of AcrIIA4 did not affect the expression of EGFP, confirming that the EGFP reduction is independent of the DNA-binding ability of Cas9 (Supplementary Figure 3B, C). To further confirm that the EGFP reduction was induced by translational repression, we tested whether the SpCas9-mediated EGFP regulation could be achieved by mRNA transfection^18,19^. By co-transfecting *SpCas9*-coding and EGFP-OFF-switch-coding mRNAs, the EGFP expression was efficiently repressed (Supplementary Figure 4). Collectively from the results, we concluded that a SpCas9-responsive translational OFF switch was designed by introducing Sp_gRNA into the 5’-UTR of mRNA.

**Figure 1.**
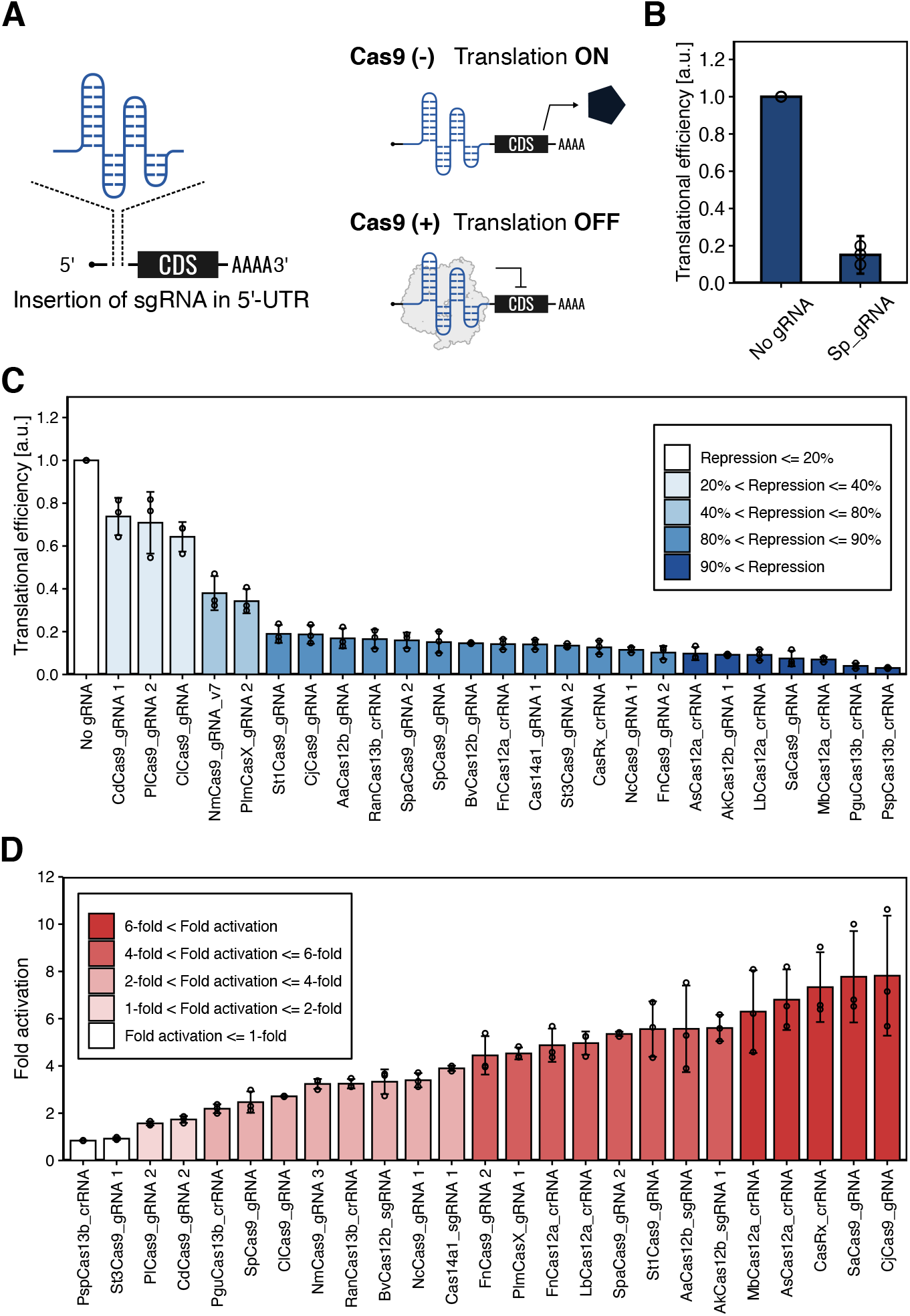
Cas proteins serve as mammalian translational modulators. (A) Design of Cas-responsive mRNA switches. sgRNAs or crRNAs are used as protein binding motifs for specific Cas proteins and inserted in the 5’-UTR of the mRNAs. The Cas protein binds to the sgRNA or crRNA region and inhibits the translation of mRNA switches. CDS: coding sequence. (B) Demonstration of SpCas9-responsive EGFP OFF switch (Sp_gRNA) as a first trial. The translational efficiency of the switch is shown. Data were normalized to the value of the control switch without the gRNA sequence (No gRNA). (C) Translational efficiencies of 25 Cas-responsive mRNA switches. Among the tested OFF switches of each Cas protein, the one with the strongest translational repression is shown. Data were normalized to the No gRNA control. See Supplementary Figure 2 for the data of switches not shown here. (D) Fold activation of the reporter expression levels in representative Cas-responsive mRNA ON switches. Among the tested ON switches of each Cas protein, the one with the strongest translational activation is shown. Data are represented as the mean ± SD from three independent experiments.

Several types of Cas proteins have been identified from diverse species^24,33,34^. We investigated whether our strategy to design OFF switches could be applied to other Cas proteins. We designed 41 different types of OFF switches that target 25 distinct Cas proteins and analyzed their responses to cognate Cas proteins (Supplementary Figure 2). Interestingly, most of the designed OFF switches repressed reporter EGFP expression in the presence of the trigger plasmid (Figure 1C). Twenty-two out of the 41 switches efficiently repressed the EGFP expression over 80% (for 20 Cas proteins), and 7 switches achieved over 90% repression. This efficiency was comparable to reported RBP (e.g., L7Ae, MS2CP and PP7CP)-responsive OFF switches that have been used to design post-transcriptional circuits^18,19,21^ (Supplementary Figure 2 and Supplementary Table 1). Although most of the designed switches efficiently responded to target Cas proteins to repress translation by just inserting reported sgRNAs or crRNAs, some switches (e.g., NmCas9-responsive switch) showed a weak response. To check the effect of the inserted sequences of sgRNA, we designed NmCas9-responsive switches by altering the sgRNA (Supplementary Figure 5). The results suggested that the repression efficiency of Cas-responsive switches can be improved by validating the sgRNA sequence. Taken together, we found that 25 Cas proteins could serve as OFF switches to regulate gene expressions post-transcriptionally.

Some Cas proteins (e.g. Cas12a and Cas13) have endoribonucleolytic activities and cleave pre-crRNAs to generate mature crRNAs^35,36^. Thus, the EGFP repression caused by Cas12a and Cas13b (Figure 1C) may be achieved by their mRNA cleaving activities. A previous paper also reported a post-transcriptional gene repression system based on a pre-crRNA processing Cas protein, Csy4, further suggesting that the repression is dependent on Csy4 RNase activity^37^. To examine the dependency on RNase activity of our OFF switches, we compared the translational repression activity of AsCas12a and its mutant (H800A), which lacks pre-crRNA processing ability^38,39^. Notably, AsCas12a (H800A) efficiently repressed EGFP expression (81% repression), although the repression efficiency was slightly weaker than that of wild-type AsCas12a (90% repression) (Supplementary Figure 6). This observation indicates that in the case of AsCas12a, RNase activity was not the main effect for the observed repression. Following these findings, we concluded that diverse Cas proteins, including those without RNase activities, can be repurposed as translational repressors.

In order to expand the versatility of Cas-mediated translational regulatory switches, we next designed Cas protein-responsive translational activator ON switches. We designed an RNA-inverter module that converts translational OFF switches to ON switches triggered by Cas proteins (Supplementary Figure 7A) by targeting mRNA degradation using an engineered non-sense mediated mRNA decay (NMD) system^27^. We found that a large majority of Cas proteins (30 out of 40 tested Cas-responsive switches showed more than 2-fold activation) functioned as triggers for translational ON switches (Figure 1D and Supplementary Figure 7B). Although the repression efficiency of the PspCas13b-responsive OFF switch was strong, its performance as an ON switch was the weakest among the Cas proteins analyzed, suggesting that the RNase activity of Cas13b may encumber the application to ON switches (Supplementary Figure 8A). Indeed, we found that the performance of ON switches was correlated with that of OFF switches for Cas proteins except for PspCas13b and PguCas13b (Supplementary Figure 8B; *r* = 0.64). This finding agrees with the mechanism of the ON switch, in which RBPs protect mRNA degradation to facilitate IRES-mediated translation^27^. In summary, we show that a wide variety of Cas proteins can serve as both translational repressors and activators through the massive construction of translational ON and OFF switches. Additionally, considering that Cas proteins are well-studied proteins for genome editing, our strategy could be combined with CRISPR-Cas associated technologies. Thus, we termed our methodology, “CaRTRIDGE”: <u>Ca</u>s-<u>R</u>esponsive <u>T</u>ranslational <u>R</u>egulation Integratable into <u>D</u>iverse <u>G</u>enomic <u>E</u>ngineering.

### Repurposing CRISPR technologies for translational regulation

Next, we aimed to conditionally regulate translation by repurposing CRISPR technologies. Conditional translational regulation with inducers (e.g. small molecules) is useful for fine-tuning gene expressions. Recent efforts have developed drug-responsive translational regulators through RBP engineering^40,41^. However, it has remained a challenge to develop various translational regulatory systems, because doing so requires additional characterization and further engineering of the individual modulators. Our methodology to regulate translation could potentially be combined with broad engineering approaches dedicated to controlling CRISPR-Cas activity, such as the drug-^42^ and light-^43^ inducible dimerization of split-Cas9. Thus, we expected available engineered CRISPR-Cas systems for genome editing and transcriptional regulation could easily be repurposed as translational regulation systems with CaRTRIDGE.

To demonstrate this possibility, we exploited the split-Cas9 system to develop a conditional translational regulation system. Previous studies reported that SpCas9 can be split into two fragments, residues 1-713 and 714-1368. The divided Cas9 protein fragments can auto-assemble and recover the genome editing ability in living cells^42,44^. Using this split-SpCas9, we constructed a translational NAND gate to repress translation only when both SpCas9 fragments are expressed (Supplementary Figure 9). Because the repression efficiency of the auto-assembled SpCas9 was low, we fused split inteins^45,46^ with the split-SpCas9 fragments^44^ to improve the performance (Figure 2A and B). The addition of inteins enhanced the translational repression comparable to wild-type SpCas9 and realized clear NAND-like behavior.

**Figure 2.**
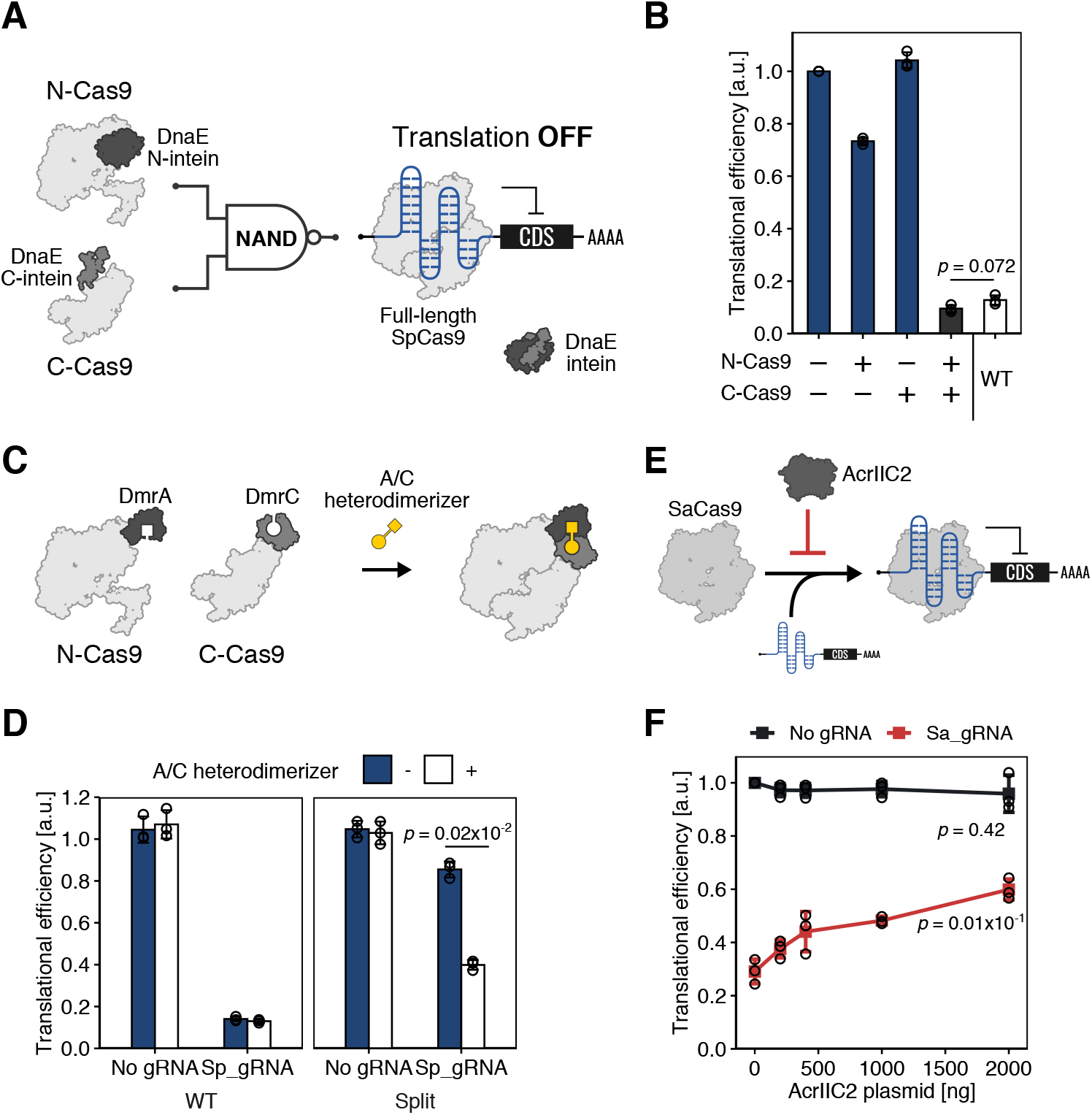
Repurposing CRISPR technologies for translational regulation. (A-B) The split-Cas9-based NAND gate. (A) The schematic architecture of NAND gate with intein-fused split-Cas9. Translational repression occurs only when both protein fragments exist. (B) The result of the reporter assay. The production of the output protein is repressed when both input proteins exist ([+, +]). The translational efficiencies were normalized to the [-, -] state. (C-D) Drug-inducible translational regulation with split-Cas9. (C) The schematic architecture. Split-Cas9s were fused with the iDimerize system. (D) The result of the reporter assay. Split-Cas9 represses translation in the presence of 500 nM A/C heterodimerizer, while wild-type Cas9 (WT) constitutively represses translation of the switch. The translational efficiencies were normalized to the control condition, in which No gRNA and No trigger (the control trigger) plasmids were co-transfected. (E-F) Translational regulation with an anti-CRISPR protein. (E) The schematic architecture of AcrIIC2-mediated translational regulation. AcrIIC2 can inhibit binding between SaCas9 protein and its sgRNA. (F) Translational efficiencies of the No gRNA control or SaCas9-responsive switch (Sa_gRNA) co-transfected with increasing amounts of the AcrIIC2-expression plasmid. The translational efficiencies were normalized by the value at 0 ng AcrIIC. *P*-values show a comparison of 0 ng and 2000 ng. Data are represented as the mean ± SD from three independent experiments. Statistical analyses were carried out with unpaired two-tailed Student’s *t*-test.

To develop a chemically controllable, SpCas9-mediated translational regulation system, we combined the split-SpCas9 with the iDimerize Inducible Heterodimer System^47^, in which separated protein fragments can be assembled by a small molecule (A/C Heterodimerizer). In the presence of A/C Heterodimerizer, the split-SpCas9 fragments were assembled and restored translational repression ability. Although the repression efficiencies were not as high as that of wild-type SpCas9, translation from the target mRNA with Sp_gRNA was significantly reduced only in the presence of the small molecule (Figure 2C, D). Altogether, these data showed that our methodology could be used to construct versatile translational regulation systems, including split-Cas proteins and small molecule-responsive systems, with existing Cas9 engineering methodologies.

In addition to engineered Cas protein-based systems, natural inhibitors of Cas proteins have been employed to fine-tune the activity of CRISPR-Cas^48,49^. For instance, Acr proteins are well-studied and promising controllers for Cas protein activity. AcrIIC2 can regulate the genome editing ability of Cas protein by inhibiting the binding between Cas proteins and their corresponding gRNAs^50^. We expected that this feature can be applied to CaRTRIDGE, because AcrIIC2 can block the interaction between Cas proteins and the corresponding RNA switches (Figure 2E). To demonstrate this, we examined the effect of AcrIIC2 on Cas9-mediated translational repression by expressing SaCas9 and a SaCas9-responsive OFF switch (Sa_gRNA-EGFP-OFF switch) with or without AcrIIC2. The translational efficiencies were increased as the amount of AcrIIC2-expression plasmid was increased (Figure 2F), indicating the inhibitory effect of the SaCas9-sgRNA interaction by AcrIIC2 and derepression of translation. Thus, Acr proteins can be used as a translational controller, and CaRTRIDGE can be combined with a variety of artificial and natural CRISPR regulatory systems.

### Evaluation of the orthogonality of Cas-responsive switches

An expanded repertoire of Cas proteins-based translational regulators should provide a means to scale-up synthetic post-transcriptional networks. Orthogonality (specificity in the interaction between the RNA and protein) is one of the key criteria of genetic circuitry when building complex circuits. To investigate the reliability of Cas-responsive OFF switches in complex circuits, we verified the orthogonality of 25 Cas-responsive switches. We transfected the switches into HEK293FT cells with each combination of switch plasmid and trigger plasmid and screened for switches that have minimal crosstalk with noncognate proteins. Among those tested, 13 Cas proteins showed high orthogonality to repress the translation of target mRNAs (Figure 3 and Supplementary Figure 10). The same tendency was observed when inverted ON switches were tested with these Cas proteins (Supplementary Figures 11 and 12). These results indicate that we successfully expanded the repertory of composable translational circuitry with orthogonal Cas protein modules. A previous study reported St1Cas9 and SaCas9 can perform genome editing orthogonally^51^. However, this pair showed crosstalk in our results, suggesting that the interaction between a Cas9 protein and its gRNA may not be the only factor for genome editing ability.

**Figure 3.**
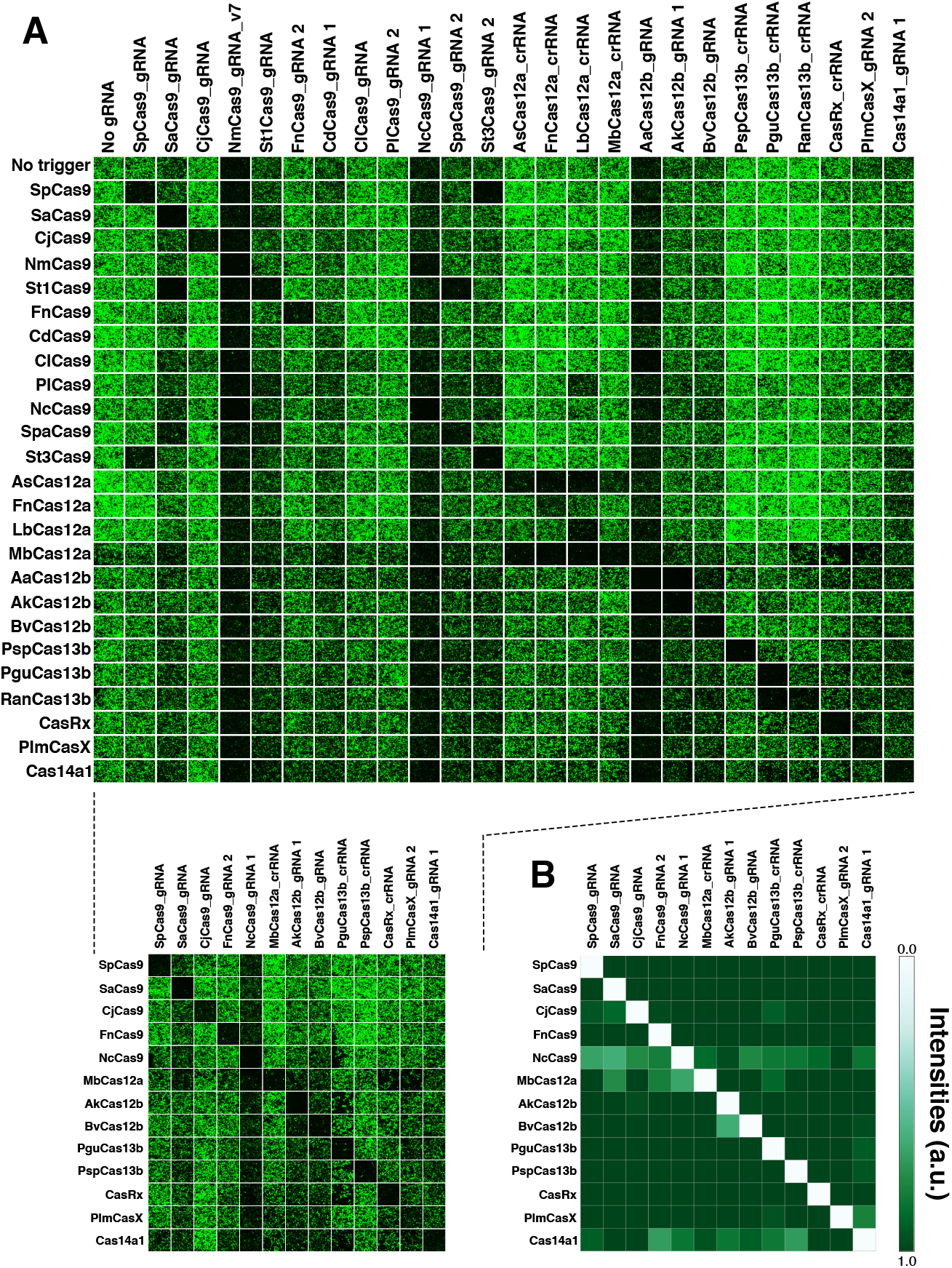
Orthogonality among representative Cas-responsive OFF switches. (A) Fluorescent cell images of a 25 × 25 orthogonality matrix of representative Cas proteins and Cas-responsive switches (top). Mutually orthogonal pairs are shown (bottom). Pruned images on the bottom indicate mutually orthogonal sets. (B) Estimated crosstalk between mutually orthogonal pairs. Data were obtained from imaging analysis. The heatmap shows the mean values from three independent experiments performed on different days.

### Dual regulations by a single Cas protein

Most transcriptional-regulatory circuits require independent triggers for transcriptional activation and repression. In contrast, one of the advantages of CaRTRIDGE is that the same trigger can serve as both a repressor (Figure 1C and Supplementary Figure 2) and an activator (Figure 1D and Supplementary Figure 7). This feature of the translational regulatory system may compact the size of the gene circuits. A recent study demonstrated that a trigger in translational regulation can target multiple translational switches and has “fan-out” ability^52^. Therefore, it is possible to regulate translational repression and activation simultaneously by a single Cas protein. To verify this point, we co-transfected translational OFF and ON switches designed to respond to SaCas9 with a trigger and the reference plasmids and observed that CaRTRIDGE controlled the translational activation and repression in a fan-out manner (Supplementary Figure 13).

Cas proteins regulate transcription by fusing transcription factors like VPR^53^ or KRAB^54^. Thus, CaRTRIDGE should enable the simultaneous control of transcription and translation by activating transcription while repressing translation. To achieve this, we used defective SpCas9 (dSpCas9) fused with a transcriptional activator, VPR, as a trigger protein (dSpCas9-VPR). This fusion protein can activate the transcription of hmAG1 guided by co-transfected Tet-responsive elements (TRE)-targeting gRNA (transcription ON) while decreasing the translation via an SpCas9-responsive OFF switch (encodes TagRFP as a reporter: referred to as Sp_gRNA-RFP) (translation OFF) (Figure 4). There was no significant difference between the hmAG1 intensity in the absence and presence of Sp_gRNA-RFP and a slight difference between the RFP intensity in the absence and presence of TRE-targeting gRNA, indicating that gRNA and Sp_gRNA-RFP have little competition for dSpCas9-VPR. Thus, CaRTRIDGE can realize dual-functional regulation (*i.e*. both translational and transcriptional layers).

**Figure 4.**
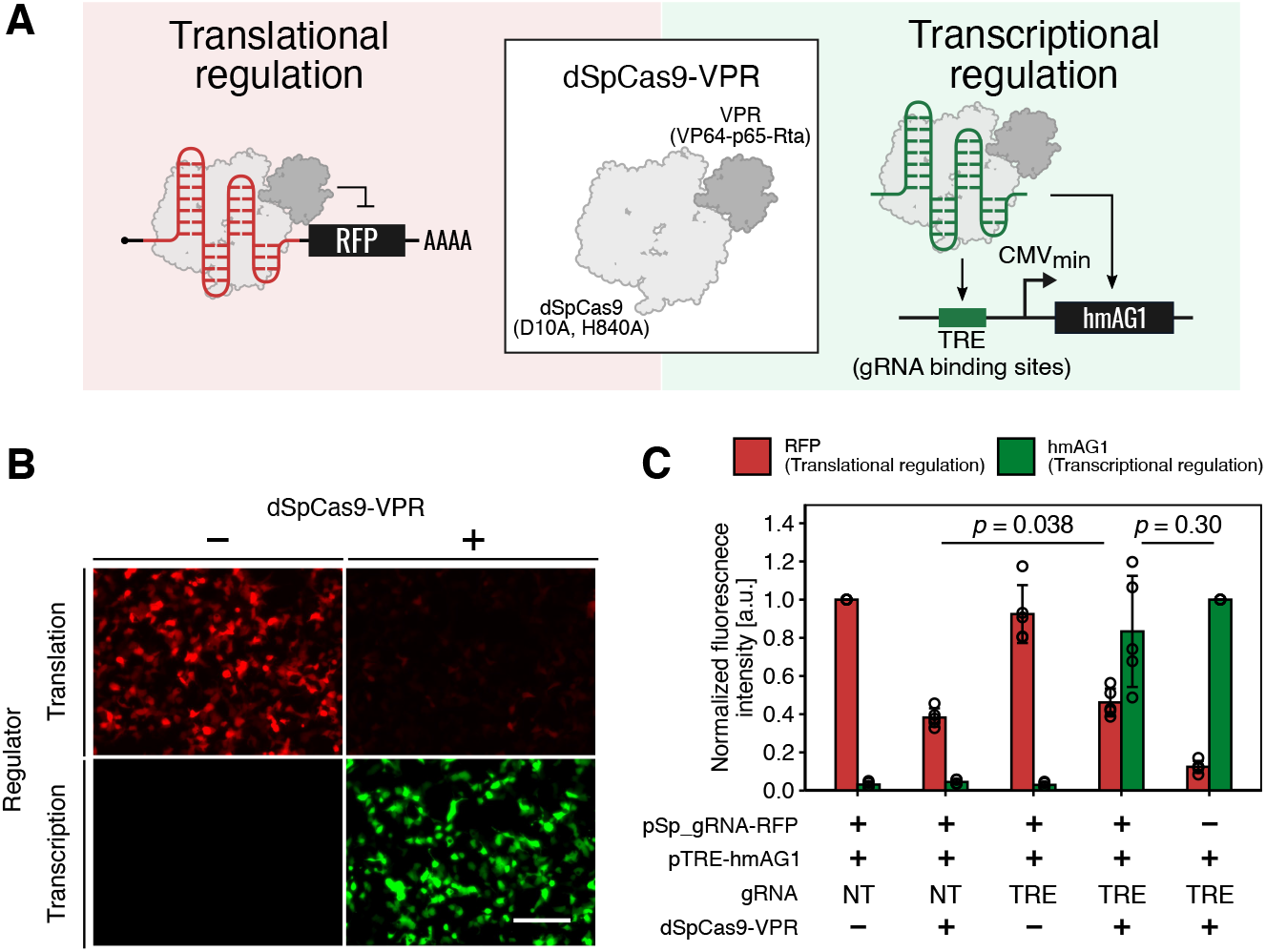
Simultaneous regulation of transcription and translation with transcription-factor fused, catalytically-dead SpCas9. (A) Schematic diagram of the simultaneous regulation system. dSpCas9-VPR (catalytically dead SpCas9 fused with a transcriptional activator) was used as the trigger protein. Translational repression and transcriptional activation were monitored with TagRFP and hmAG1, respectively. Tet-responsive elements (TRE) were used as the gRNA binding site. (B) Fluorescence microscopy images in each condition. Translational repression and transcriptional activation were observed simultaneously when dSpCas9-VPR was transfected. Scale bar, 200 μm. (C) Quantitative data of reporter expression levels. The data were obtained by imaging analysis. TagBFP was co-transfected as a reference and used to define the transfection-positive area. pSp_gRNA-TagRFP: plasmid for expressing TagRFP, whose translation is regulated by SpCas9. pTRE-hmAG1: plasmid for expressing hmAG1, whose transcription is regulated by Tet-responsive promoter. TRE: TRE-targeting gRNA. NT: non-targeting gRNA. Error bars represent the mean ± SD from 5 independent experiments. Statistical analyses were carried out with unpaired two-tailed Student’s *t*-test.

### Validating the composability of Cas-responsive RNA switches

To verify the composability of Cas-responsive OFF switches in multi-layered translational circuits, we explored the AND gate circuit as a representative 2-input multi-layered logic circuit (Figure 5). An AND circuit expresses the output only in the presence of both inputs (input pattern [11]). Two input translational AND circuits can be constructed using 3 OFF switches, as shown in Figure 5A. We arbitrarily chose 6 Cas-responsive switches and constructed mediator plasmids that express Cas protein “C” except when both Cas protein “A” and “B” are present. Accordingly, we could prepare and test 60 patterns of AND circuits that were composed of different combinations of 3 of 6 Cas-responsive switches, which is the largest number of translational AND gates ever tested (Figure 5B, 5C and Supplementary Table 2). To measure the similarity between the intended function of the AND gate and experimental data, we calculated the net fold-change (ratio of the mean fluorescence intensity of ON state ([11] state) and OFF states ([00], [10], and [01])) and the cosine similarity, which were recently proposed as metrics to evaluate genetic circuits^13^. Twenty-one circuits showed angles less than or equal to 20°, which is comparable to previously constructed translational AND circuits^18^ (Figure 5D). All circuits that included the NcCas9-responsive switch showed angles over 20°, which might be due to the low basal expression level of the mediator proteins and insufficient repression for the reporter (Figure 5C, 5D and Supplementary Table 2).

**Figure 5.**
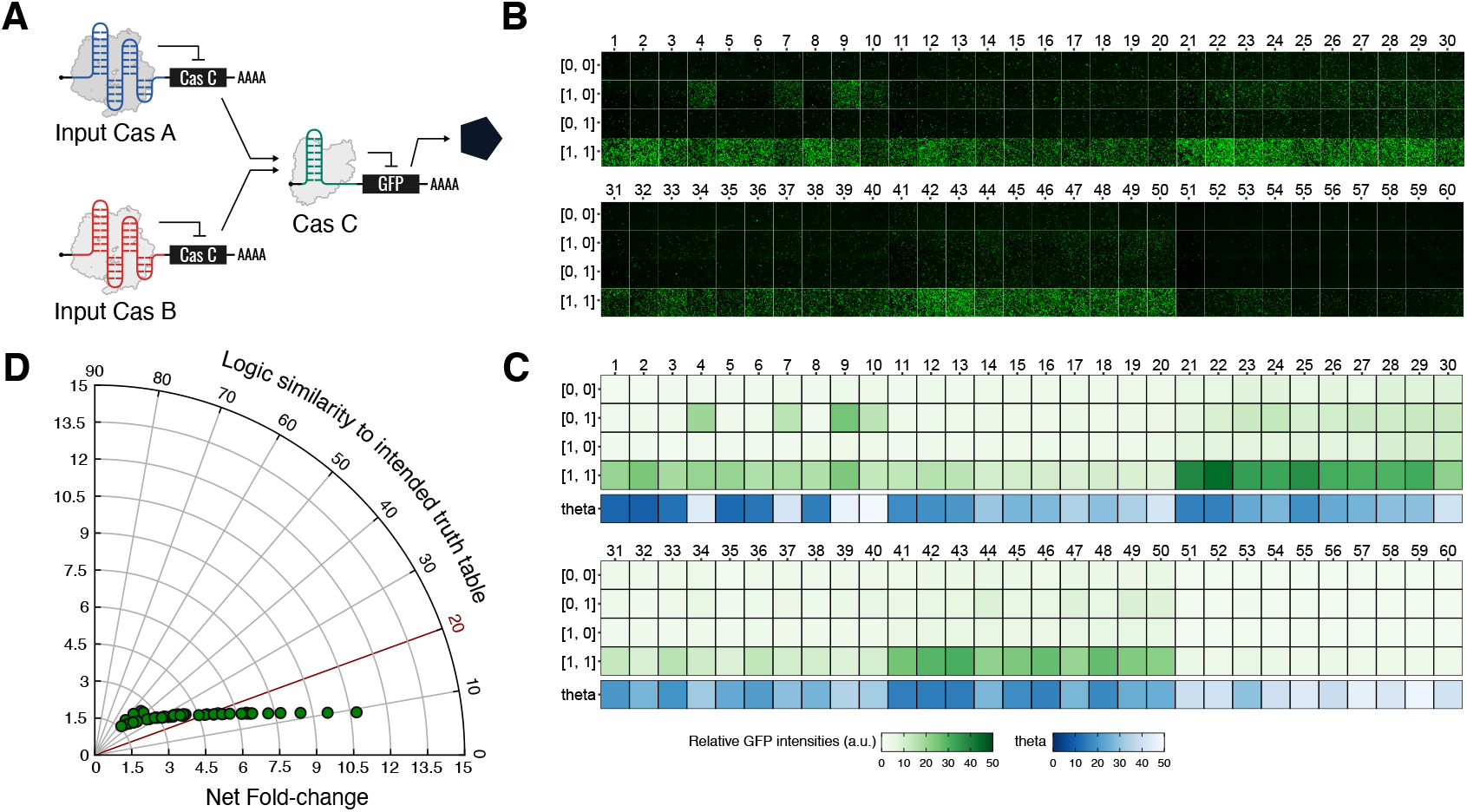
Multi-layered translational circuits with Cas-responsive switches. (A-D) Combinatorial validation of Cas-responsive switches’ composability. Sixty distinct 2-input translational AND gate circuits were tested. (A) The schematic diagram of a 2-input AND gate composed of three Cas-responsive switches. Six Cas proteins were selected and arranged as Cas protein A, B (as inputs), and C (as a mediator). Thus, a total of 60 circuits were tested. (B) Fluorescence microscopy images of the 60 distinct 2-input AND gates with four input patterns. The absence and presence of the inputs are indicated by 0 and 1, respectively. The ideal AND gate shows reporter expression only in the [1, 1] state. (C) Heatmap of the GFP expression level and vector proximity angles. The data were obtained by flow cytometry analysis. (D) A polar coordinate plot of the net fold-change and vector proximity angles. Data are represented as the mean from three independent experiments.

### An arithmetic circuit via compacted modules

Our CaRTRIDGE methodology showed high orthogonality, dual-functionality, and composability in complex circuits. These results encouraged us to construct higher-order synthetic networks with complex signal wiring using Cas proteins and corresponding gRNAs. Previous transcription-translation coupled binary arithmetic circuits required 4 genetic modules, 2 transcriptional controllers and 2 translational controllers^5^. We considered building a binary arithmetic circuit with fewer modules, because CaRTRIDGE can simultaneously control transcription and translation using a single Cas protein. We designed a half-subtractor with two catalytically dead Cas proteins, dSpCas9 and dSaCas9, fused with VPR as inputs (Figure 6). Both proteins are used as transcriptional regulators and are orthogonal to each other. To monitor the calculated difference (D) and the borrow (B_0_), we chose the fluorescent proteins hmAG1 and TagBFP as the output signals. dSaCas9-VPR and dSpCas9-VPR activated the transcription of Sp_gRNA-TagBFP and Sa_gRNA-TagBFP mRNA, respectively, and the translation of these mRNAs was orthogonally suppressed by dSpCas9-VPR and dSaCas9-VPR. TagBFP expression is therefore regulated in an XOR manner (outputs are produced only when exactly one Cas protein is inputted (Figure 6A, TagBFP). Meanwhile, the expression of hmAG1 is regulated in a “dSpCas9 NIMPLY dSaCas9” manner: hmAG1 is produced only when dSpCas9 alone is present. To achieve this logic, the gRNA is quarried out from the 3’-UTR of the Sa_gRNA-TagBFP transcript by a hammerhead ribozyme (HHR) and a hepatitis delta virus ribozyme (HDVR) and guides dSpCas9-VPR to the promoter region of hmAG1^55,56^. We also inserted a MALAT1 triplex upstream of the ribozymes to stabilize the transcript after loss of the poly-A tail^57^. The 5’-UTR of the hmAG1 transcript contains SaCas9 gRNA, which is translationally regulated by dSaCas9. Thus, dSpCas9 activates the transcription of hmAG1, but its translation is suppressed by dSaCas9 (Figure 6A, hmAG1). Imaging analysis indicated the designed half-subtractor behaved as expected (Figure 6B). Thus, CaRTRIDGE can implement complex logic operations with more compactness and stage programmability in mammalian cells compared with other synthetic circuits.

**Figure 6.**
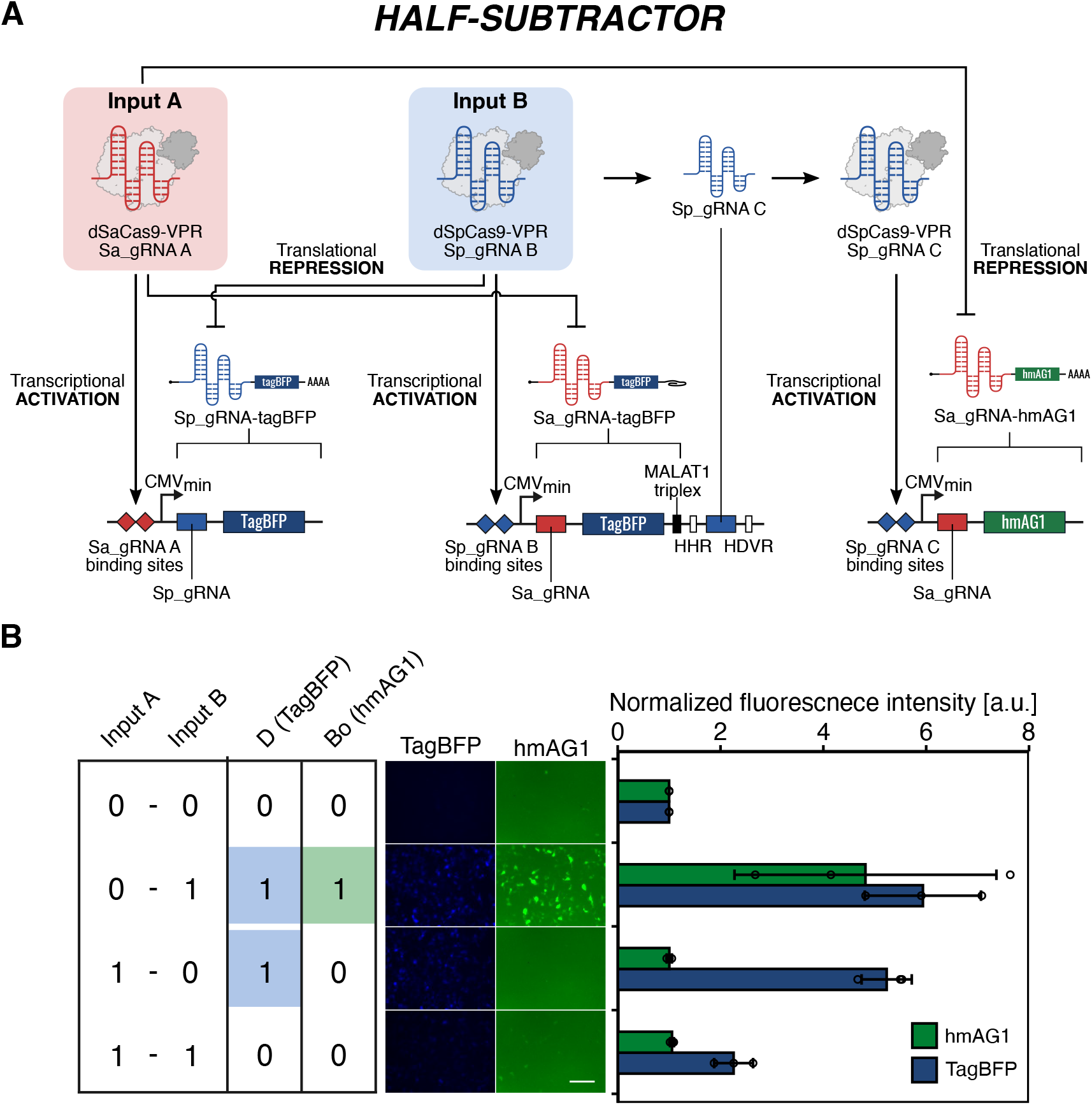
Design and processing performance of a half-subtractor with fewer bio-modules. (A) Schematic architecture of the 2-input half-subtractor. VPR-fused dSpCas9 and dSaCas9 are used as the initial inputs (top left). The transcription of mRNA switches (middle) from the corresponding plasmids (bottom). The translation is regulated by the indicated input dCas9. (B) Fluorescence microscopy images and analysis of the half-subtractor performing arithmetic subtraction of the SpCas9 input from the SaCas9 input by calculating the difference, D (TagBFP), and borrow, B_0_ (hmAG1). Data are represented as the mean ± SD from three independent experiments. Scale bar, 200 μm.

## Discussion

In this study, we developed CaRTRIDGE to repurpose Cas proteins into translational repressors and activators. We initially tested 25 Cas proteins for their translational repression efficiency and found 20 switches showed over 80% repression. Among these, 14 switches were constructed by just inserting reported gRNA at the 5’-UTR, and the other 6 switches were developed by validating the gRNA sequences. We converted these Cas-responsive OFF switches to ON switches by an RNA-inverter strategy^27^ and generated 21 types of ON switches. Furthermore, we designed 26 different orthogonal regulators (13 OFF and ON switches each), 60 translational logic gates,and constructed a complex arithmetic circuit via compacted modules. To our knowledge, this is the largest number of translational regulatory devices and translational logic gates ever tested, since only a limited number of translational regulatory proteins have been used to build post-transcriptional circuits. Importantly, our CaRTRIDGE strategy, in which Cas proteins are used as translational modulators, is compatible with other conventional CRISPR technologies. For example, the use of split-Cas9 and anti-CRISPR proteins enabled us to perform small molecule-based and conditional translational regulation, suggesting that transcriptional and genome regulatory systems for CRISPR can be combined with CaRTRIDGE. Thus, CaRTRIDGE will provide promising modules for synthetic gene circuits.

Recent efforts have expanded the catalog of available synthetic parts for constructing gene circuits. The recombinase-based approach BLADE^13^ provides highly robust circuits. However, it is an irreversible operation, and it is difficult to reset the cell state. COMET^11^, which uses ZF-TFs as the transcriptional regulator, and another similar approach that uses meganuclease^14^ can perform reversible regulation in principle. These transcriptional regulation strategies, however, cannot be applied to DNA-free transgene methods like mRNA delivery. In contrast, CaRTRIDGE can potentially drive synthetic circuits at the translation level with mRNA transfection (Supplementary Figure 4). Furthermore, CaRTRIDGE is compatible with transcriptional circuits so that more complex genetic circuits can be built with fewer bio-modules (Figure 6). Recent studies have constructed post-translational circuits by exploiting proteases^16^, coiled-coil domains^15^, and de novo designed heterodimeric domains^58^. Given that CaRTRIDGE is an RBP-based regulatory system, these engineering approaches can be incorporated to make even more complex circuits. Indeed, one study has already tested Cas9 with such engineering^59^.

During this research, several studies reported post-transcriptional regulation using Cas proteins. Among Cas proteins, two studies used only Cas12a and mainly focused on how to supply crRNA from mRNA, which encodes Cas12a itself^38,60^. These studies tested several Cas12a variants, but most have strong crosstalk^61^ (Figure 3), which hinders large-scale connections of bio-modules. Another study reported a novel post-transcriptional-regulation system, PERSIST, which uses Cas proteins that have endoribonuclease activity^62^. This system greatly expanded strategies for synthetic post-transcriptional regulation. However, the tested Cas proteins were limited to only those with RNase activity. That is, mRNA degradation by endonucleases is essential to make switches with PERSIST, whereas CaRTRIDGE does not have this restriction. Another study also reported a translational activation system with SpCas9^63^, but limited its application to yeast and required to seek the optimal position of the gRNA insertion. In contrast, we validated the properties of far more Cas proteins (25 Cas proteins investigated as translational regulators) in mammalian cells, suggesting the potential power of CaRTRIDGE and its compatibility with CRISPR technologies.

Overall, CaRTRIDGE has several advantages for the design and construction of gene circuits. (1) As more CRISPR-associated genes are discovered due to huge efforts in the genome editing field, we can exploit more and more Cas proteins as translational modulators. (2) CaRTRIDGE can be combined with existing and even undeveloped CRISPR technology. In addition to three translational-regulation methods demonstrated in this study (Figure 2; split-Cas9, drug-inducible regulation, and Acr), other possible approaches include light-inducible regulation^43^, protease-combined regulation^59^, CRISPR-based gene circuits^6^, and molecular event recorders^10^. These conditional regulations will allow us to finely tune the transgene expression at the post-transcriptional level. (3) CaRTRIDGE enables the simultaneous regulation of transcription and translation by a single Cas protein (Figure 4), reducing the verbosity of the circuits. This advantage can potentially achieve higher information density and processing power with fewer bio-modules (Figure 6). Therefore, CaRTRIDGE will provide “biological Integrated Circuits” and facilitate the development of easily designable biocomputers. For the same feature, metabolic burden in the chassis will likely be decreased^64^. (4) CaRTRIDGE provides the opportunity to study the design principle of protein-responsive mRNA OFF and ON switches. In this study, we developed a library of Cas-responsive switches with a wide variety of translational modulators. Further elucidation of the relationship between translation efficiency and biochemical and biophysical parameters (e.g. structural features and binding kinetics) may lead to improving mRNA switches. (5) CaRTRIDGE will support new therapeutic modalities, such as mRNA-based medicine, which conducts fine tuning of the transgene expression at the post-transcriptional level, and cell therapy, which requires precise control of cellular behaviors that cannot be addressed by the simple expression of transgenes.

In addition, CaRTRIDGE may contribute to the genome editing field. For example, our system provides an initial screening to select interactable Cas proteins-sgRNA pairs in human cells. Cas9s with inefficient translational repression (e.g. ClCas9-ClCas9_gRNA, FnCas9-FnCas9_gRNA 1) seem to lack genome editing ability in mammalian cells^65,66^. CaRTRIDGE can be used to eliminate Cas proteins-gRNA pairs that are inactive in cells. Furthermore, some small molecules are reported to disrupt SpCas9-DNA interactions and control SpCas9 genome editing ability^67^. CaRTRIDGE may facilitate the discovery of these inhibitors against Cas protein-gRNA complexes, which could provide powerful genome editing regulators.

One of the major drawbacks of CaRTRIDGE is the gene size of the Cas protein and it may prevent some applications. In the present study, we showed the applicability of split-Cas9 to reduce the size of the plasmid (Figure 2), but a more efficient gene delivery system free of size limitations should be developed. Another solution is the use of smaller Cas proteins. Our data suggest that controlling translation is independent of the Cas protein size (Supplementary Table 1). Thus, CaRTRIDGE will directly benefit from the discovery of novel Cas proteins^68,69^. Because several RBPs used in synthetic mRNA switches can control translation naturally^70^, we anticipate that some Cas proteins may control translation by directly interacting with mRNA in prokaryotes. In conclusion, we believe that CaRTRIDGE will inspire not only RNA- and RBP-based medicine and cell therapies, but also future genome editing technologies and CRISPR studies.

## Author contribution

S.K. and H.S. managed the project. S.K. conceived the idea. M.H. found the translational repression ability of SpCas9. S.K., H.O., M.H. and H.S. designed the project. S.K., H.O., M.H., and T.K. performed the experiments and analyzed the data. H.O. mainly prepared the figures under the supervision of S.K. S.K., H.O., M.H. and H.S. wrote the manuscript. S.K., H.O. and M.H. contributed equally to this work.

## Acknowledgments

We thank Dr. Peter Karagiannis (Kyoto University) and Dr. Zoher Gueroui (École Normale Supérieure) for critical reading of the manuscript, Yusuke Shiba (Kyoto University) for helping with the experiments, and Miho Nishimura and Yuko Kono for their administrative support. We also thank Dr. Hideyuki Nakanishi (Tokyo Medical and Dental University) for the intein information, Yoshihiko Fujita (Kyoto University) for establishing the imaging quantification method, and Dr. Akitsu Hotta (Kyoto University) for providing the AsCas12a ORF. This work was supported by JSPS KAKENHI Grant Number JP 19K16110 (to S.K.), 19J21199 (to H.O.), 20K15777 (to M.H.), 15H05722, and 20H05626 (to H.S.).

## Methods

### Construction of plasmids

#### Switch plasmids

pAptamerCassette-EGFP (available at Addgene, #140288) was digested by AgeI and BamHI. Single-stranded oligo DNAs were annealed to generate double-stranded DNA and then ligated with the digested plasmid. Links to sequences of all the generated switch plasmids are listed in Supplementary Table 3.

#### Trigger plasmids

The open reading flame (ORF) of trigger proteins was amplified by PCR using appropriate primers. The amplicons were digested by restriction enzymes and then inserted downstream of the CMV promoter of pcDNA3.1-myc-HisA (invitrogen). Links to sequences of all the generated trigger plasmids are listed in Supplementary Table 3.

#### Split-Cas9 plasmids

To construct split-Cas9, hCas9 (obtained from Addgene, #41815) was used as the template. N_Cas9 and C_Cas9 were amplified by inverse PCR using appropriate primers. To generate drug-responsive Cas9, DmrA and DmrC were amplified and then fused to each split-Cas9 (N_Cas9 and C_Cas9) using the In-Fusion HD Cloning Kit (Clontech). Links to sequences of all the generated plasmids are listed in Supplementary Table 3.

#### Plasmid for multiple genetic circuits

The ORF of each Cas protein was amplified using appropriate primers. Using each switch plasmid as a template, the backbone of the plasmids was amplified by inverse PCR. The ORF was inserted in the plasmid backbone using the In-Fusion HD Cloning Kit. Links to sequences of all the generated plasmids were listed in Supplementary Table 3.

All Plasmids were purified using the Midiprep kit (QIAGEN or Promega)

### Construction of IVT (*in vitro* transcription) template

The ORF of SpCas9 (for trigger mRNA) and iRFP670 (for reference mRNA), 5’-UTR fragment, and 3’-UTR fragments were amplified by PCR using appropriate primers and template plasmids or oligo DNAs.

The IVT templates of trigger mRNA and reference mRNA were generated by fusion PCR using the 5’-UTR fragment, the 3’-UTR fragment, the ORF, and appropriate primers.

To make an IVT template of switch mRNA, a fragment including the 5’-UTR and ORF was amplified from SpCas9-responsive switch plasmid by PCR using appropriate primers. The fragment and the 3’-UTR fragment were fused by fusion PCR using appropriate primers.

To remove the template plasmids in the PCR products, 1 µL DpnI was added to each reaction, and the mixture was incubated at 37°C for 30 min. All PCR products were purified using the MinElute PCR purification kit (QIAGEN) or Monarch PCR & DNA Cleanup Kit (NEB) according to the manufacturer’s protocols. The set of primers is listed in Supplementary Table 4.

### mRNA synthesis and purification

All mRNAs were prepared using a MEGAscript T7 Transcription Kit (Thermo Fisher Scientific). To reduce the immune response, pseudouridine-5′-triphosphate (ΨTP) and 5-methylcytidine-5′-triphosphate (m5CTP) (both from TriLink BioTechnologies) were used instead of natural UTP and CTP, respectively, except for switch mRNA. The IVT reaction included 1 × Enzyme mix, 1 × Reaction buffer, 7.5 mM ΨTP or UTP, 7.5 mM m5CTP or CTP, 7.5 mM ATP, 1.5 mM GTP, 6 mM Anti Reverse Cap Analog (TriLink BioTechnologies), and the template DNA. The mixture was incubated at 37°C for 6 h. To remove the template DNA, TURBO DNase (Thermo Fisher Scientific) was added to the mixture and incubated at 37°C for 30 min. Then the reaction mixtures were purified using a FavorPrep Blood/Cultured Cells total RNA extraction column (Favorgen Biotech) or Monarch RNA Cleanup kit

(New England Biolabs) and incubated with Antarctic Phosphatase (New England Biolabs) at 37°C for 30 min. The reaction mixtures were purified again using the RNeasy MinElute Cleanup Kit (QIAGEN) or Monarch RNA Cleanup kit according to the manufacturer’s protocols.

All IVT mRNA sequences are shown in the Supplementary Sequences.

### Cell culture

HEK293FT cells (Invitrogen) were cultured in DMEM High glucose (nacalai tesque) supplemented with 10% FBS (Biosera), 2 mM L-Glutamine (Invitrogen), 0.1 mM Non-Essential Amino Acids (Invitrogen) and 1 mM Sodium Pyruvate (Sigma) at 37°C with 5% CO_2_ in a humidified cell culture incubator. In Supplementary Figure 13, HEK293FT cells that constitutively express TagBFP were used. The cells were cultured in the same manner as HEK293FT cells.

### Plasmid transfection

HEK293FT cells were seeded in 24-well (1.0×10^5^ cells/well), 96-well (2.0×10^4^ cells/well), or 384-well (2.5×10^3^ cells/well) plates. Appropriate plasmids were transfected into the cells using Lipofectamine 2000 (Invitrogen) according to the manufacturer’s protocols. Details of the transfection conditions for each experiment are shown in Supplementary Table 5. iRFP670 plasmid was used as a transfection control in all experiments except for those shown in Figure 4, for which TagBFP plasmid was used. For the experiments shown in Figure 2C and D, A/C heterodimerizer-containing media (final concentration 500 nM) were prepared and replaced before the transfection.

### mRNA transfection

HEK293FT cells were seeded in 24-well plates (1.0×10^5^ cells/well). Appropriate mRNAs were transfected into the cells using Lipofectamine MessengerMAX (Invitrogen) according to the manufacturer’s protocols.

Details of the transfection conditions for each experiment are shown in Supplementary Table 5

### Cell imaging

Before the flow cytometry measurements, cell images were captured using the Cytell Cell Imaging System (GE Healthcare Life Sciences). To capture each fluorescent image, the following channels were used: Blue channel (Ex 390 nm / Em 430 nm) for TagBFP, Green channel (Ex 473 nm / Em 512.5 nm) for EGFP and hmAG1, Orange channel (Ex 544 nm / Em 588 nm) for TagRFP, and Red channel (Ex 631 nm / Em 702 nm) for iRFP670. The captured images were analyzed using ImageJ (NIH).

### Imaging analysis and calculations

The tiff files of the captured fluorescent cell images were analyzed with ImageJ. The background intensities were subtracted using the rolling ball algorithm/method before calculation. The cell area was defined by a reference (iRFP670 or, for experiments shown in Figure 4, TagBFP) positive place. The median value of the reporter/reference within the defined area was calculated and used for the analysis.

In the orthogonality heatmaps, intensities were calculated using the following formulas:

Defined value (DV) = median value of the reporter/reference in each cell.

Relative value (RV) = (DV of trigger +) / (DV of trigger −).

Intensity = (RV) / (RV of No gRNA sample).

### Flow cytometry measurements

The cells were washed with PBS, treated with 0.25% Trypsin-EDTA (Thermo Fisher Scientific) and incubated at 37°C with 5% CO_2_ for 5 min. The cells were analyzed using an Accuri C6 (BD Bioscience) flow cytometer with FL1 (533/30 nm) and FL4 (675/25 nm) filters.

### Flow cytometry data analysis

Flow cytometry data sets were analyzed using FlowJo version 10.5.3 (BD Biosciences) and Excel (Microsoft). Gates were generated using mock samples. Data from debris were eliminated when preparing forward-versus side-scatter dot plots (FSC-A versus SSC-A). Then, events on the chart edges in the dot plots of the EGFP intensity versus the iRFP670 intensity were removed. In the histogram where iRFP670-intensity is displayed on the X-axis, the iRFP670-positive (reference-positive) gate was defined (Supplementary Figure 1B). In the following analysis, the median of reporter/reference of each cell was calculated from the reference positive population using FlowJo.

Translational efficiency is defined using the following formulas:

Normalized intensity (NI) = 1000 × median of the ratio (reporter intensity/reference intensity) of each cell.

Relative intensity (RI) = (NI of trigger +) / (NI of trigger −).

Translational efficiency = (RI) / (RI of No gRNA sample).

All values were normalized by the value of the No gRNA sample.

In Supplementary Figure 2, all RI values were normalized by the value of the ON state No gRNA sample.

The fold activations in Figure 1D and Supplementary Figure 7 are defined as RI.

The fold change in Supplementary Figure 14C is defined as RI in ON switch and the reciprocal of RI in OFF switch.

### Statistical analysis

All statistical analysis was performed by unpaired two-tailed Student’s t-test using R software or Excel (Microsoft).

### Multiple Sequence Alignment for phylogenetic tree

Representative Cas9 gRNA sequences that showed crosstalk were analyzed with Multiple Sequence Comparison by Log-Expectation (MUSCLE: https://www.ebi.ac.uk/Tools/msa/muscle/). The result was exported as a FASTA file, and a phylogenetic tree was drawn by Python3.

