## Supplementary Information for "Programmable mammalian translational modulators by CRISPR-associated proteins"

1 Co-first authors, equally contributed
2 Department of Life Science Frontiers, Center for iPS Cell Research and Application, Kyoto University, 53 Kawahara-cho, Shogoin, Sakyo-ku, Kyoto 606-8507, Japan.
3 Department of Life Science Frontiers, Center for iPS Cell Research and Application, Kyoto University, 53 Kawahara-cho, Shogoin, Sakyo-ku, Kyoto 606-8507, Japan; Graduate School of Medicine, Kyoto University, Kyoto 606-8507, Japan.
4 Faculty of Medicine, Kyoto University, Kyoto 606-8507, Japan

*Corresponding author
**Lead contact

**Contents**

Supplementary Figure 1| Schematic diagrams of experimental procedures in plasmid transfection.
Supplementary Figure 2| Original validation data of each Cas-responsive switch related to Figure 1C.
Supplementary Figure 3| SpCas9-responsive reporter reduction is independent of transcriptional regulations.
Supplementary Figure 4| The performance of SpCas9-responsive switch in RNA transfection.
Supplementary Figure 5| The performances of NmCas9-responsive switches.
Supplementary Figure 6| Effect of RNase ability in AsCas12a-responsive switch
Supplementary Figure 7| Cas proteins serve as the triggers of RNA-inverters.
Supplementary Figure 8| Correlation of the performance in Cas-responsive OFF switches and ON switches
Supplementary Figure 9| The translational repression with auto-assemble Split-Cas9.
Supplementary Figure 10| Orthogonality heat-map among representative Cas-responsive OFF switches related to Figure 3A.
Supplementary Figure 11| Fluorescent cell images of 25 × 25 orthogonality matrix of representative Cas proteins and Cas-responsive ON switches.
Supplementary Figure 12| Heat-map of 25 × 25 orthogonality matrix of representative Cas proteins and Cas-responsive ON switches.
Supplementary Figure 13| Simultaneous regulation of translational activation and repression with SaCas9.
Supplementary Figure 14| Phylogenetic tree of Cas9 gRNA used in OFF switch orthogonality.
Supplementary Table 1| Information about Cas proteins used in this study.
Supplementary Table 2| Profile of the 60 AND gates in Figure 5.
Supplementary Table 3| Key plasmids used in this study.
Supplementary Table 4| Primers, template oligo DNA for generating synthetic mRNAs.
Supplementary Table 5| Transfection tables of all experiments performed in this study.
Supplementary Sequences

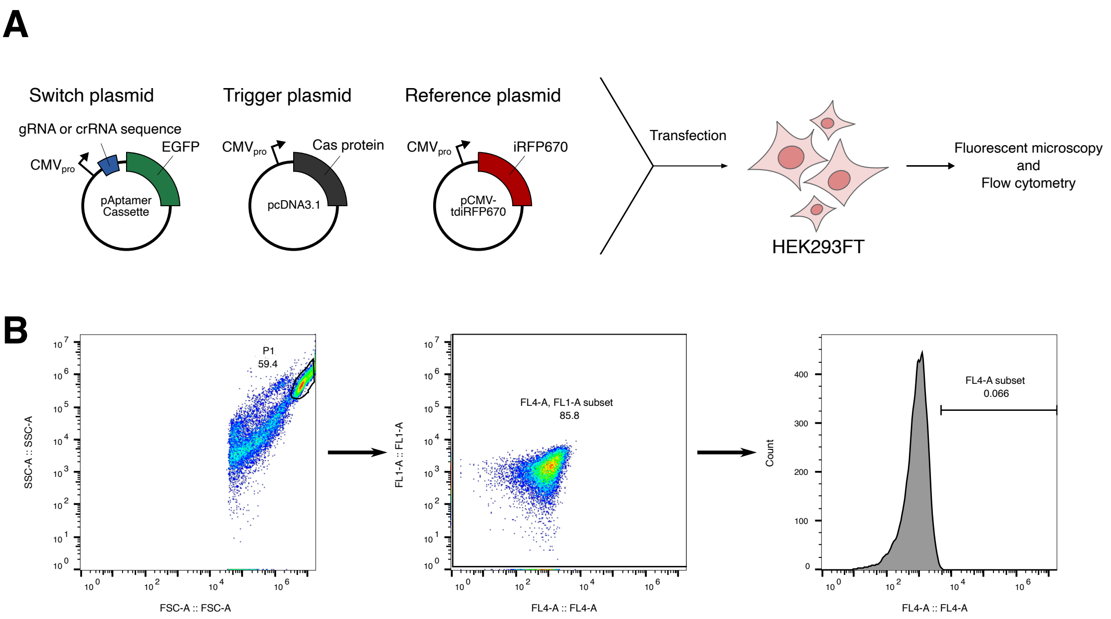

**Supplementary Figure 1|**

**Schematic diagrams of experimental procedures in plasmid transfection.**

(A) Schematic diagrams of standard procedure. Three plasmids (switch plasmid, trigger plasmid, and reference plasmid) were transfected into HEK293FT cells. The cells were analyzed by fluorescent microscopy and flow cytometer. (B) Gating procedures of flow cytometry. Flow cytometry datasets were analyzed using FlowJo version10.5.3 (see also Methods section). Live cells were gated in the forward scatter (FSC) versus side scatter (SSC) plot to eliminate debris. The remaining P1-positive events were plotted in the FL1-A (EGFP expression, Y-axis) versus FL4-A (iRFP670 expression, X-axis) and events on each axis line were ruled out by the gate (FL4-A, FL1-A subset). Then, the FL4-A subset gate was generated based on untransfected samples and thereby transfection-positive (FL4-positive) populations were defined.

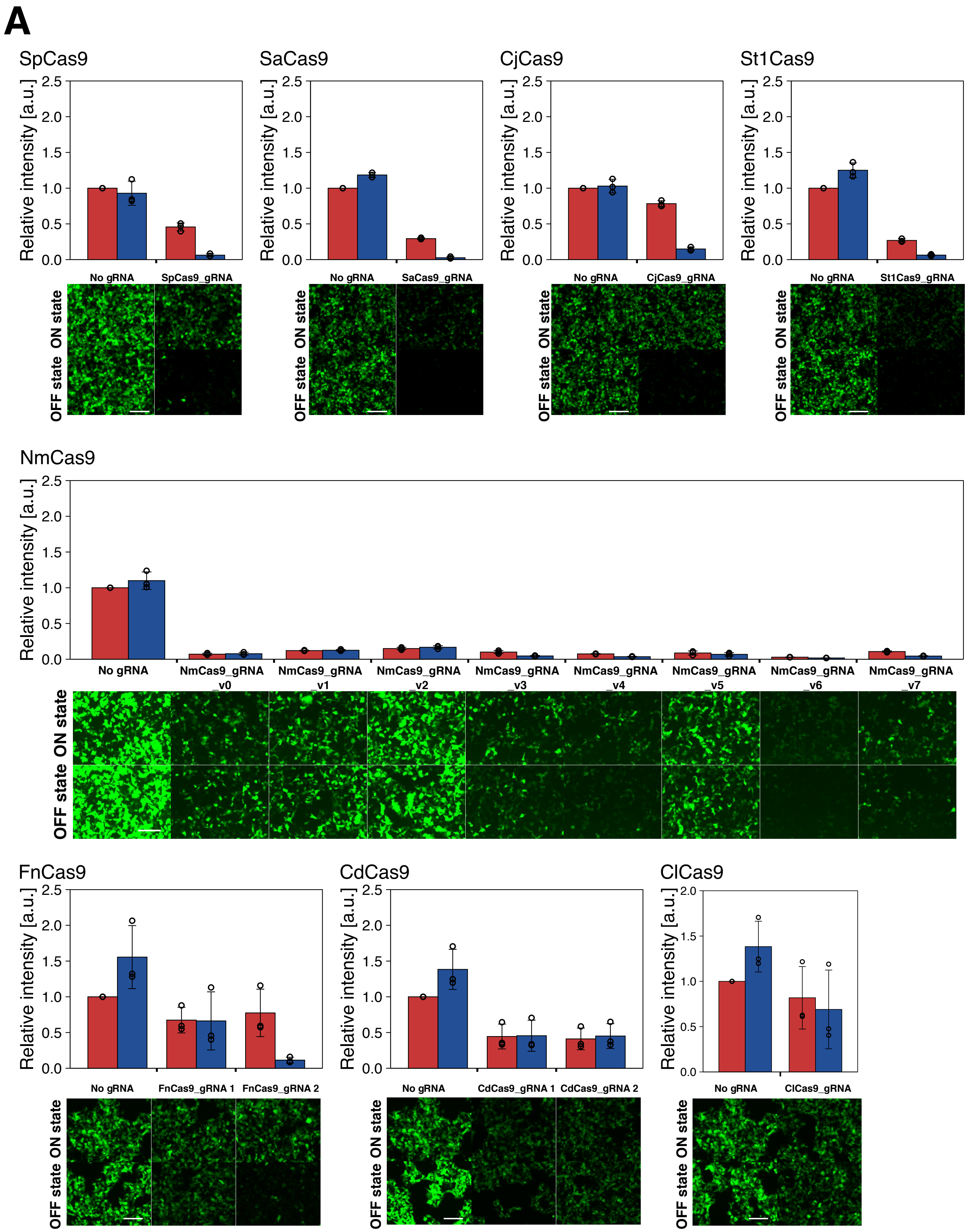
A
SpCas9
SaCas9
CjCas9
St1Cas9
Relative intensity [a.u.]
0.0
0.5
1.0
1.5
2.0
2.5
No gRNA
SpCas9_gRNA
SaCas9_gRNA
CjCas9_gRNA
St1Cas9_gRNA
ON state
OFF state
NmCas9
NmCas9_gRNA _v0
NmCas9_gRNA _v1
NmCas9_gRNA _v2
NmCas9_gRNA _v3
NmCas9_gRNA _v4
NmCas9_gRNA _v5
NmCas9_gRNA _v6
NmCas9_gRNA _v7
FnCas9
CdCas9
ClCas9
FnCas9_gRNA 1
FnCas9_gRNA 2
CdCas9_gRNA 1
CdCas9_gRNA 2
ClCas9_gRNA

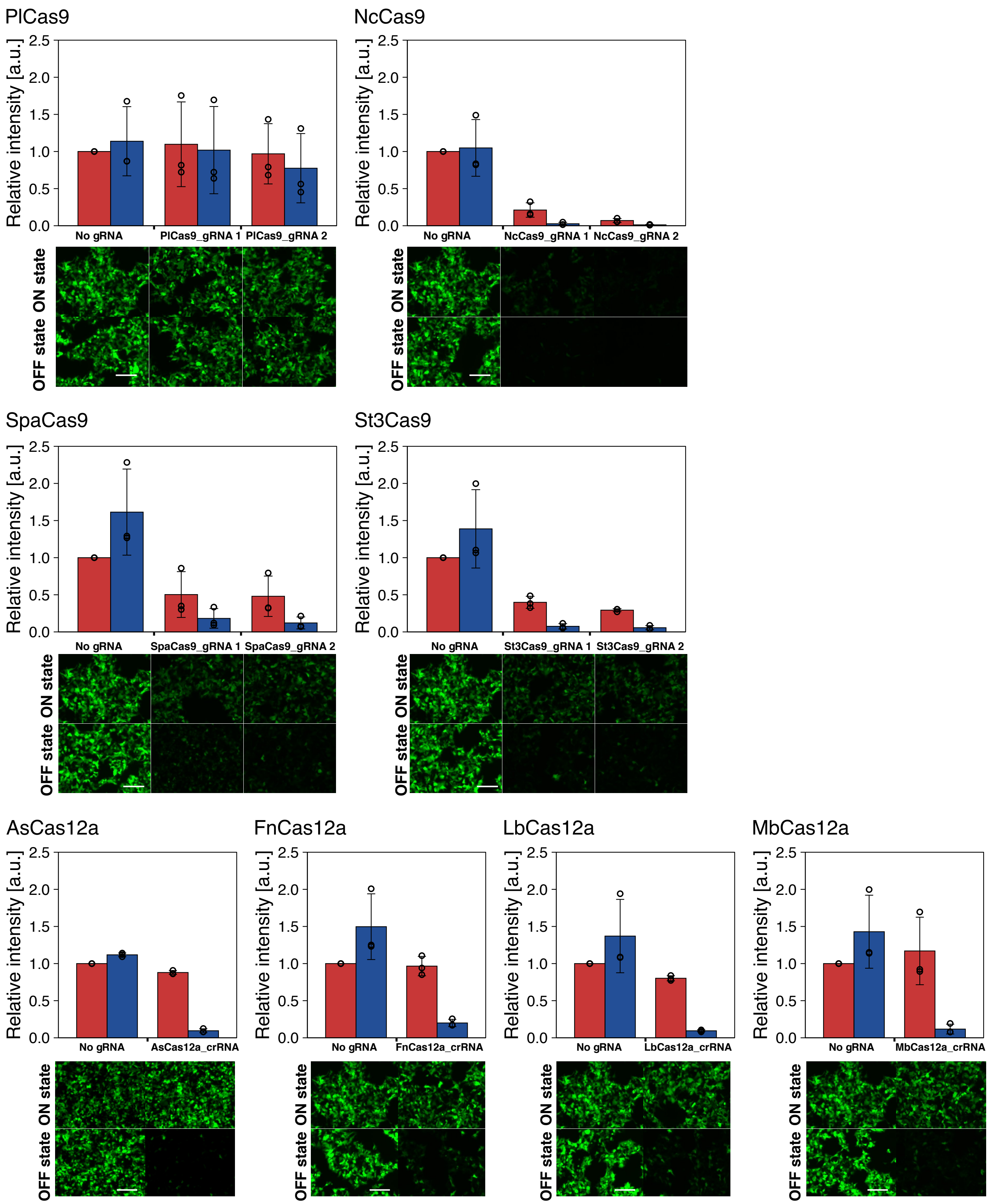
PlCas9
NcCas9
SpaCas9
St3Cas9
AsCas12a
FnCas12a
LbCas12a
MbCas12a
Relative intensity [a.u.]
0.0
0.5
1.0
1.5
2.0
2.5
No gRNA
PlCas9_gRNA 1
PlCas9_gRNA 2
NcCas9_gRNA 1
NcCas9_gRNA 2
SpaCas9_gRNA 1
SpaCas9_gRNA 2
St3Cas9_gRNA 1
St3Cas9_gRNA 2
AsCas12a_crRNA
FnCas12a_crRNA
LbCas12a_crRNA
MbCas12a_crRNA
ON state
OFF state

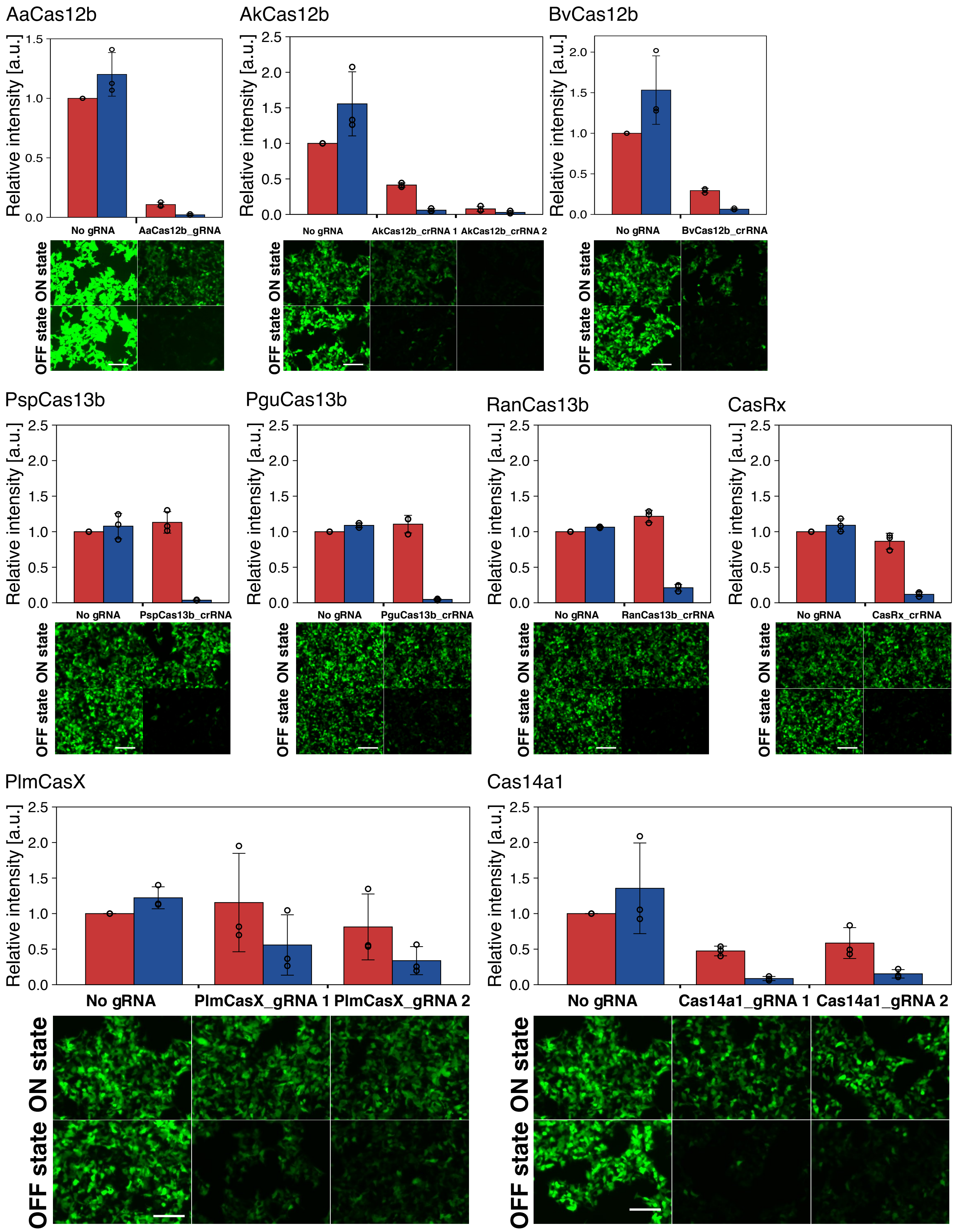
AaCas12b
Relative intensity [a.u.]
1.5
1.0
0.5
0.0
No gRNA
AaCas12b_gRNA
ON state
OFF state
AkCas12b
Relative intensity [a.u.]
2.5
2.0
1.5
1.0
0.5
0.0
No gRNA
AkCas12b_crRNA 1
AkCas12b_crRNA 2
ON state
OFF state
BvCas12b
Relative intensity [a.u.]
2.0
1.5
1.0
0.5
0.0
No gRNA
BvCas12b_crRNA
ON state
OFF state
PspCas13b
Relative intensity [a.u.]
2.5
2.0
1.5
1.0
0.5
0.0
No gRNA
PspCas13b_crRNA
ON state
OFF state
PguCas13b
Relative intensity [a.u.]
2.5
2.0
1.5
1.0
0.5
0.0
No gRNA
PguCas13b_crRNA
ON state
OFF state
RanCas13b
Relative intensity [a.u.]
2.5
2.0
1.5
1.0
0.5
0.0
No gRNA
RanCas13b_crRNA
ON state
OFF state
CasRx
Relative intensity [a.u.]
2.5
2.0
1.5
1.0
0.5
0.0
No gRNA
CasRx_crRNA
ON state
OFF state
PlmCasX
Relative intensity [a.u.]
2.5
2.0
1.5
1.0
0.5
0.0
No gRNA
PlmCasX_gRNA 1
PlmCasX_gRNA 2
ON state
OFF state
Cas14a1
Relative intensity [a.u.]
2.5
2.0
1.5
1.0
0.5
0.0
No gRNA
Cas14a1_gRNA 1
Cas14a1_gRNA 2
ON state
OFF state

**B**

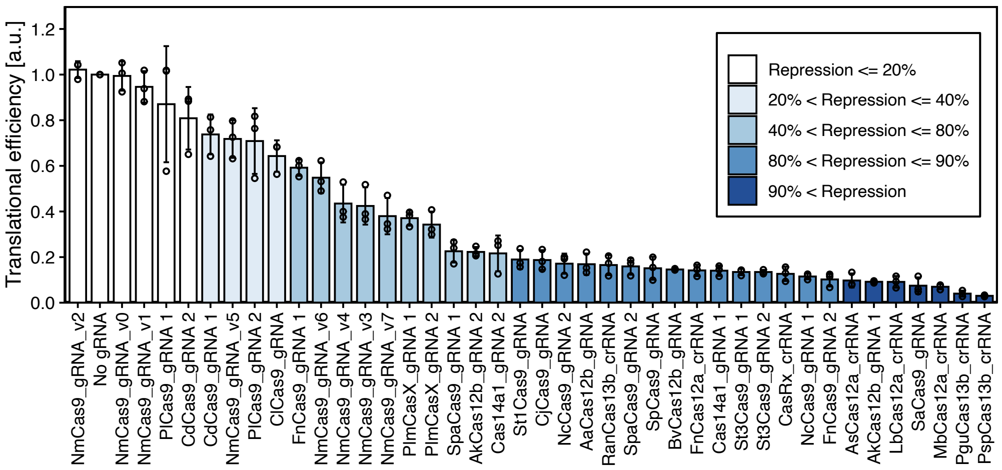

**Supplementary Figure 2|**

**Original validation data of each Cas-responsive switch related to Figure 1C.**

(A) Validated trigger Cas proteins are stated at the top. The bar charts show the calculated reporter expression levels. Error bars represent standard deviations. ON (without trigger) and OFF (with trigger) states are colored as red and blue, respectively. Scale bars in cell images represent 200 μm. (B) Translational efficiencies of the reporter expression levels in all tested Cas-responsive mRNA OFF switches. Data are represented as the mean ± SD from three independent experiments.

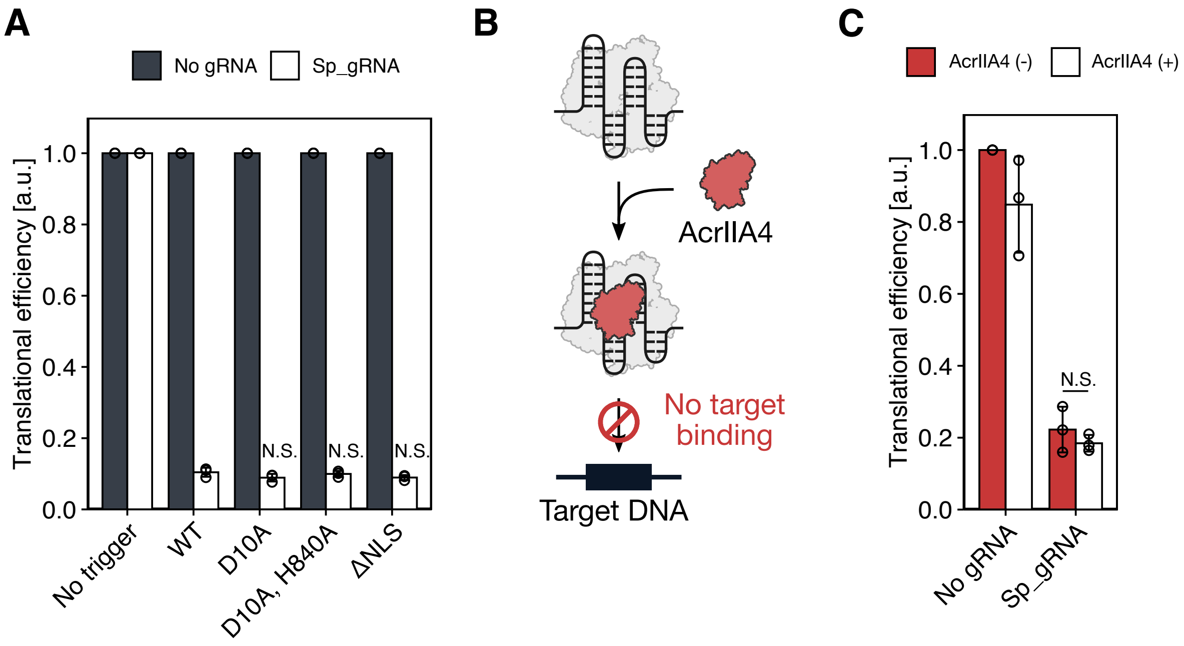

**Supplementary Figure 3|**

**SpCas9-responsive reporter reduction is independent of transcriptional regulations.**

(A) The translational efficiencies in SpCas9 mutants transfection. Each SpCas9 mutant did not affect the translational efficiency of Sp_gRNA compared with wild type (WT). (B) Schematic diagram of AcrIIA4 mediated binding inhibition between DNA and SpCas9 RNP complex. (C) The translational efficiencies in co-transfection of AcrIIA4. Error bars represent standard deviations. Statistical analyses were carried out with unpaired two-tailed Student's *t*-test. N.S., not significant ($p \geq 0.05$).

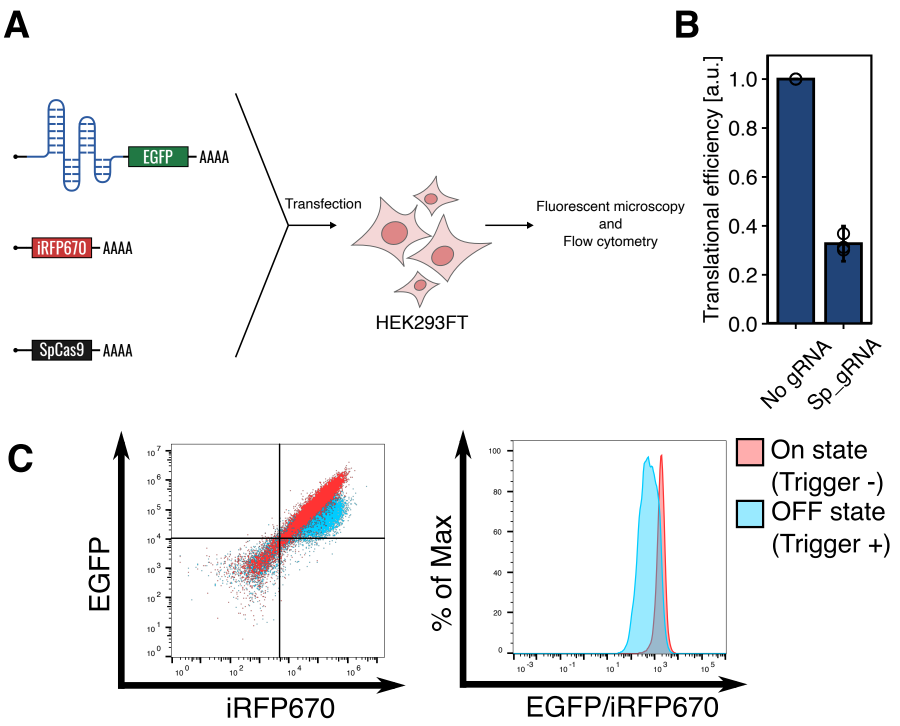

**Supplementary Figure 4|**

**The performance of SpCas9-responsive switch in RNA transfection.**

(A) Schematic diagrams of experimental procedures in RNA transfection. Three mRNAs (switch mRNA, trigger mRNA (SpCas9 mRNA), and reference mRNA (iRFP670 mRNA)) were transfected into HEK293FT cells. The cells were analyzed by fluorescent microscopy and flow cytometer. (B) The translational efficiencies in RNA transfection of SpCas9-responsive switches. Error bars represent standard deviations. (C) Representative dot plots and the histograms of expression level from switch mRNA with (blue) and without (red) SpCas9 mRNA.

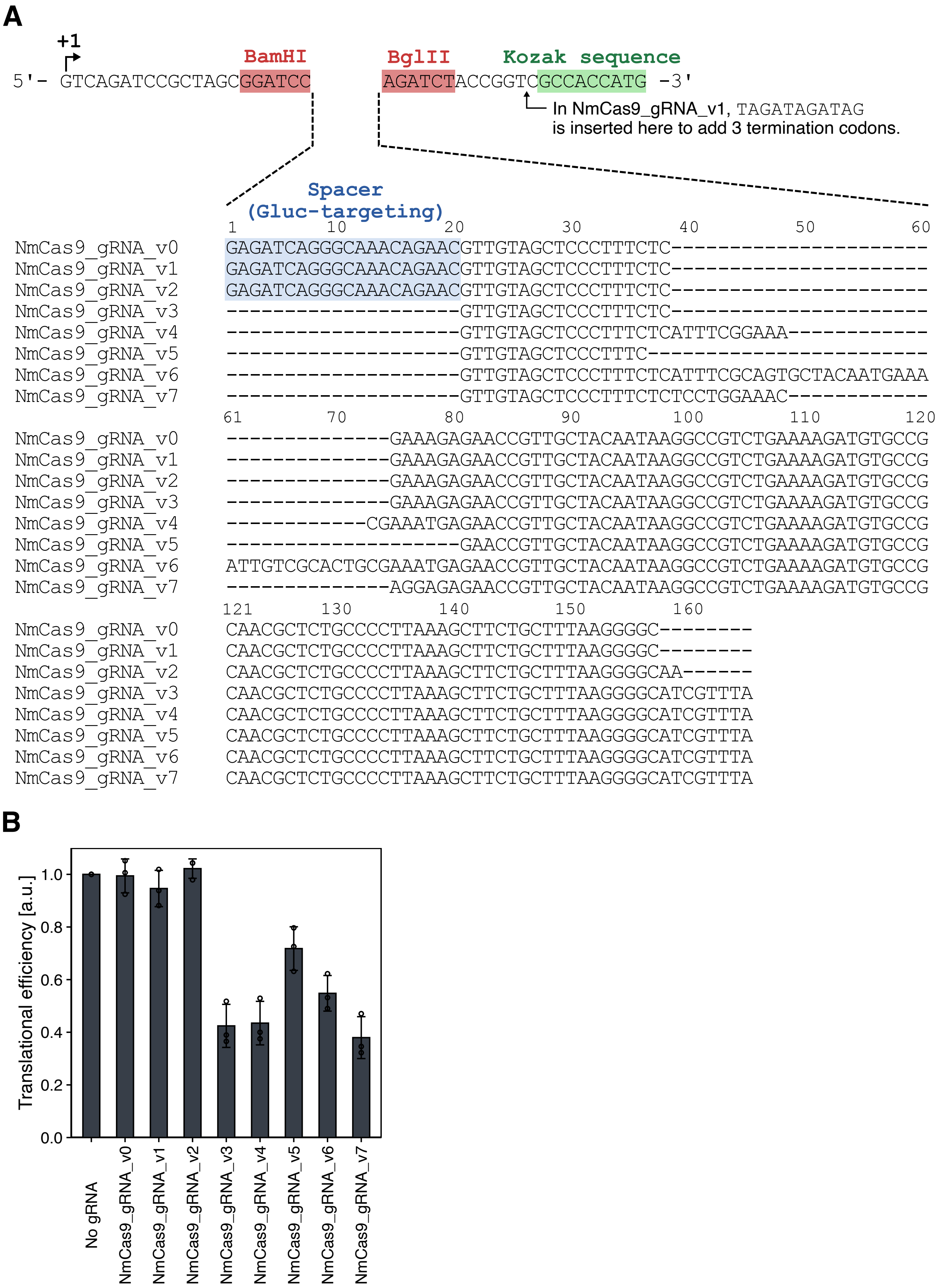

**Supplementary Figure 5|**

**The performances of NmCas9-responsive switches.**

(A) Sequence variations of NmCas9-responsive switches. The surrounding sequence of the inserted gRNA sequence is also shown. (B) The translational efficiencies of NmCas9-responsive switches. The switch plasmid with NmCas9_gRNA_v7 sequence showed the best performance and was used for the following OFF switch experiments. Error bars represent standard deviations.

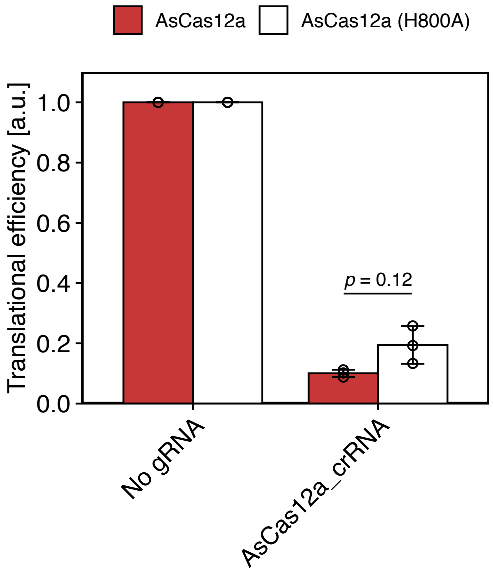

**Supplementary Figure 6|**

**Effect of RNase ability in AsCas12a-responsive switch**

Comparison of translational repression efficiency between wild type AsCas12a and its mutant (H800A). AsCas12a (H800A) lacks RNase activity but maintains other enzyme activity. Values were normalized by the value in the condition when WT and No gRNA reporter were co-transfected. Data are represented as the mean ± SD from three independent experiments. Error bars represent standard deviations. Statistical analyses were carried out with unpaired two-tailed Student's *t*-test.

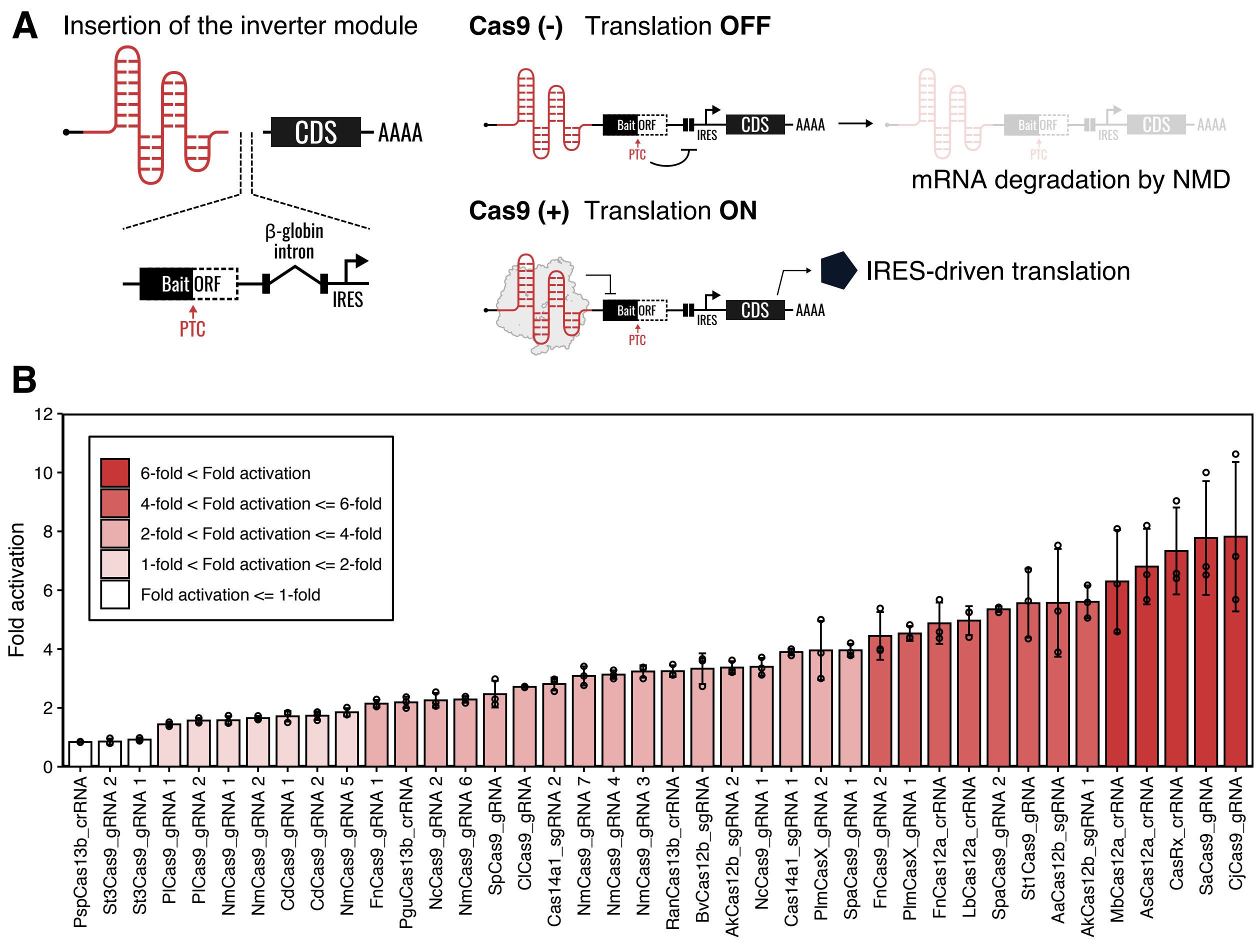

**Supplementary Figure 7|**

**Cas proteins serve as the triggers of RNA-inverters.**

(A) Design of Cas-responsive mRNA ON switches with RNA-inverter. Inverter module was inserted between the protein binding motifs for specific Cas proteins and CDS. The Cas protein binds to the sgRNA or crRNA and protects mRNA switches from the non-sense mediated mRNA decay (NMD). CDS: coding sequence. (B) Fold activations of the reporter expression levels in all tested Cas-responsive mRNA ON switches. Data are represented as the mean ± SD from three independent experiments.

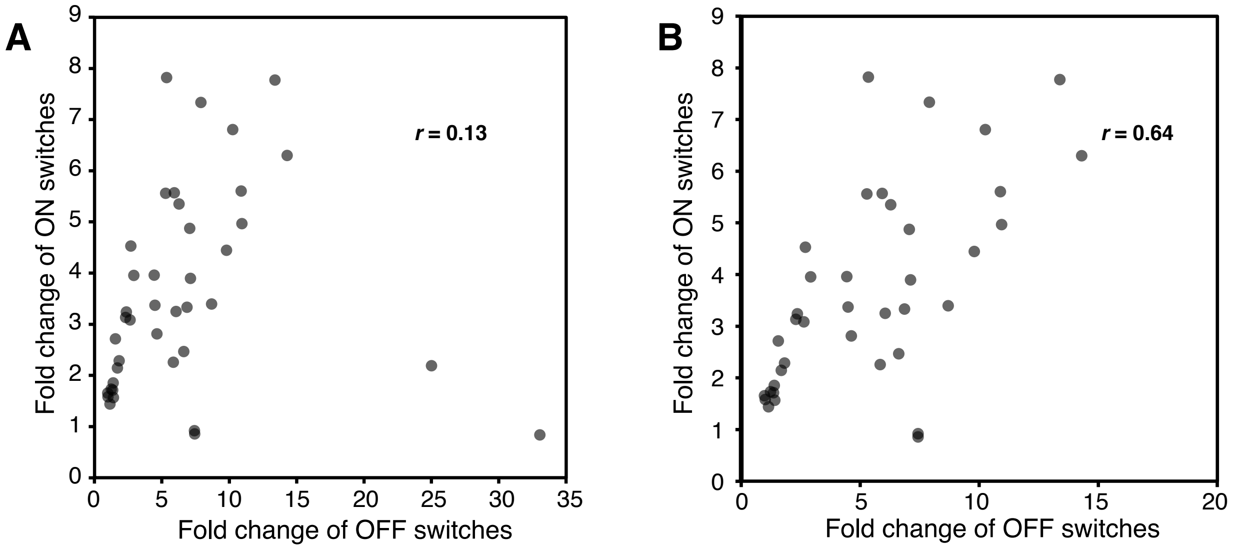

**Supplementary Figure 8|**

**Correlation of the performance in Cas-responsive OFF switches and ON switches**

(A) Comparison of all Cas-responsive OFF and ON switches shown in Supplementary Figure 2B and Supplementary Figure 7B. Fold Change of OFF switch was calculated as inverse number of translational efficiencies. (B) Comparison of Cas-responsive OFF and ON switches without PspCas13b- and PguCas13b-responsive ones. Pearson correlation coefficients (*r*) are shown in the charts.

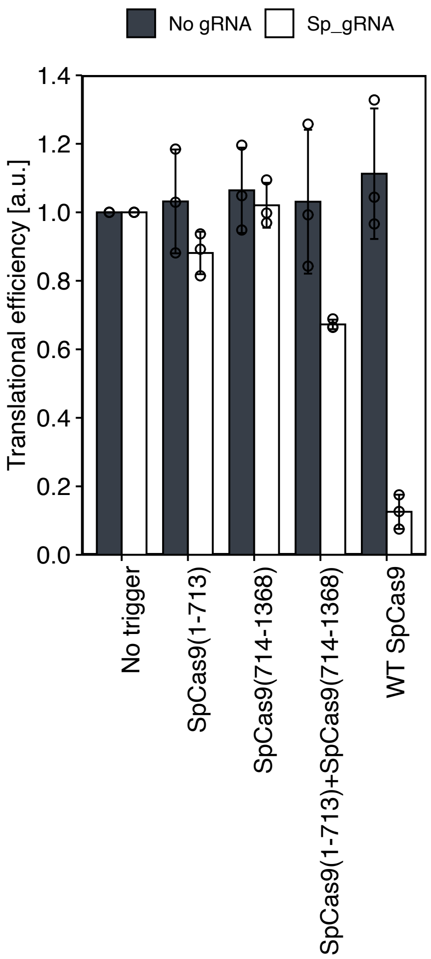

**Supplementary Figure 9|**

**The translational repression with auto-assemble split-Cas9.**

The translational repression is conducted only when both protein fragments exist. The reporter assay resulted in 33% repression when both of the input proteins exist. Data are represented as the mean ± SD from three independent experiments.

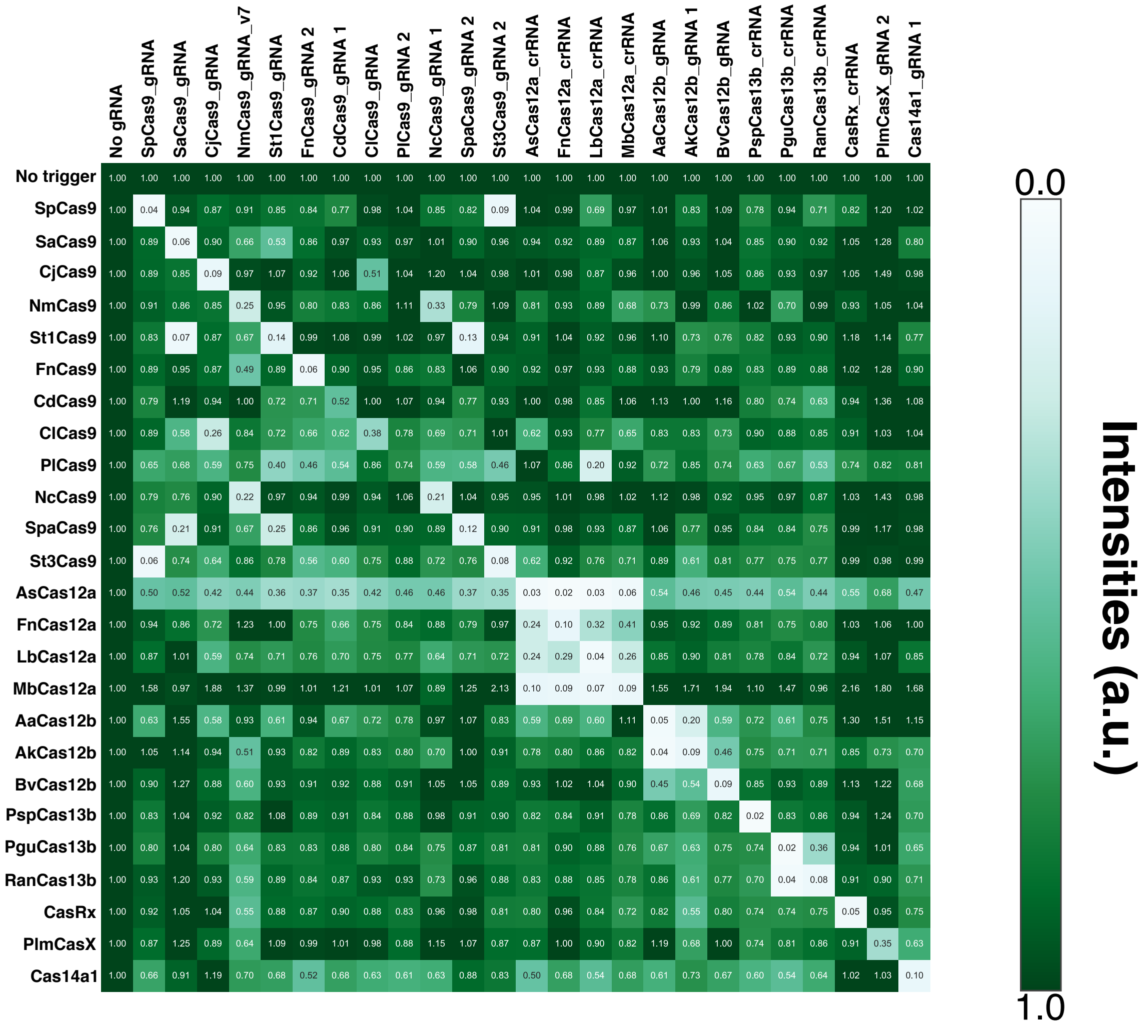

**Supplementary Figure 10|**

**Orthogonality heat-map among representative Cas-responsive OFF switches related to Figure 3A.**

Heat-map of 25 × 25 orthogonality matrix of representative Cas proteins and Cas-responsive switches with the indicated mean values from Imaging analysis.

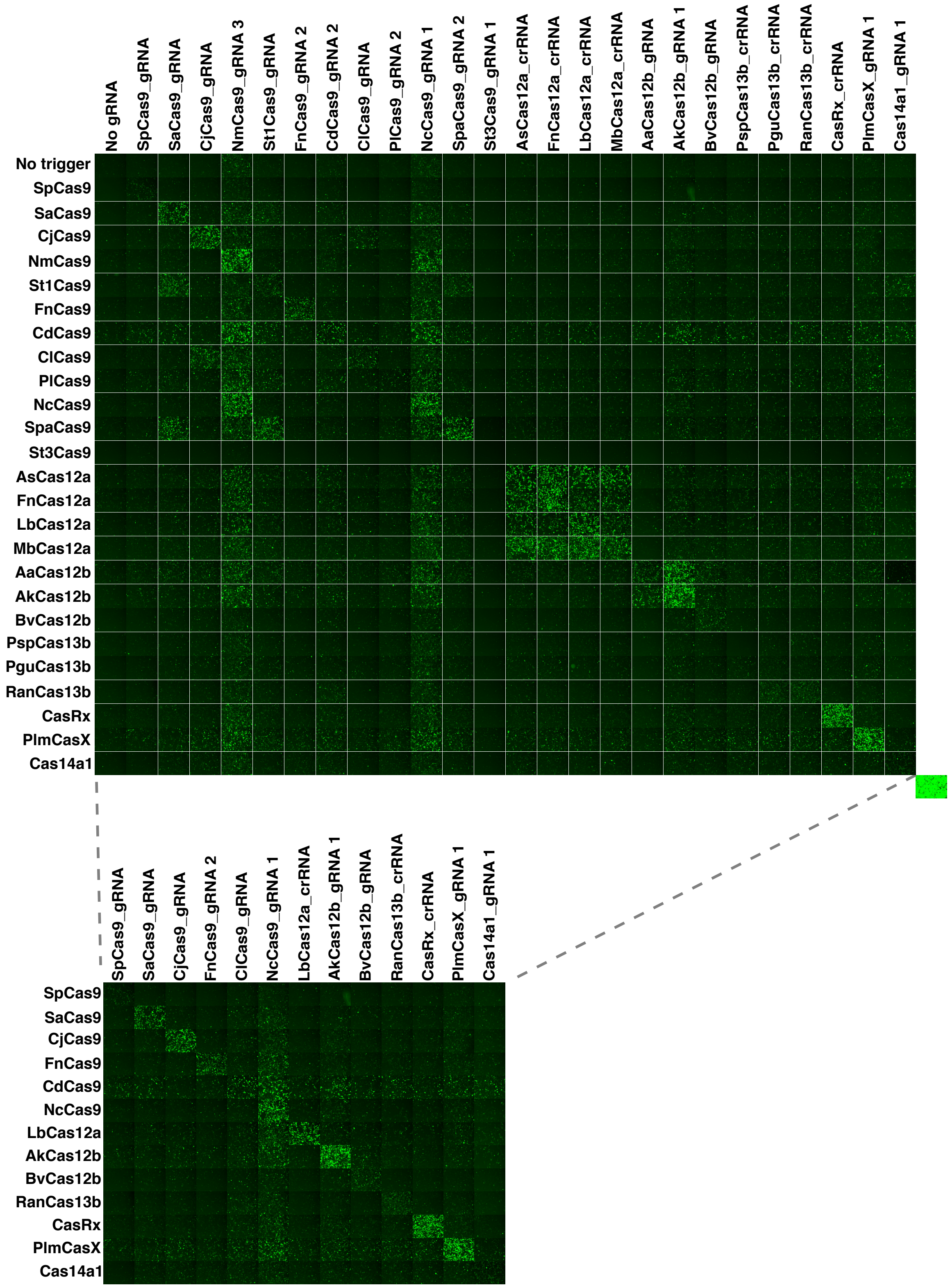

**Supplementary Figure 11|**

**Fluorescent cell images of 25 × 25 orthogonality matrix of representative Cas proteins and Cas-responsive ON switches.**

A fluorescent image of cells transfected with a plasmid constitutively expressing EGFP is shown at the bottom right of the upper panel. Pruned images in the bottom indicate mutually orthogonal sets.

| | No gRNA | SpCas9_gRNA | SaCas9_gRNA | CjCas9_gRNA | NmCas9_gRNA 3 | St1Cas9_gRNA | FnCas9_gRNA 2 | CdCas9_gRNA 2 | ClCas9_gRNA | PlCas9_gRNA 2 | NcCas9_gRNA 1 | SpaCas9_gRNA 2 | St3Cas9_gRNA 1 | AsCas12a_crRNA | FnCas12a_crRNA | LbCas12a_crRNA | MbCas12a_crRNA | AaCas12b_gRNA | AkCas12b_gRNA 1 | BvCas12b_gRNA | PspCas13b_crRNA | PguCas13b_crRNA | RanCas13b_crRNA | CasRx_crRNA | PlmCasX_gRNA 1 | Cas14a1_gRNA 1 |
|---|---|---|---|---|---|---|---|---|---|---|---|---|---|---|---|---|---|---|---|---|---|---|---|---|---|---|
| No trigger | 1.00 | 1.00 | 1.00 | 1.00 | 1.00 | 1.00 | 1.00 | 1.00 | 1.00 | 1.00 | 1.00 | 1.00 | 1.00 | 1.00 | 1.00 | 1.00 | 1.00 | 1.00 | 1.00 | 1.00 | 1.00 | 1.00 | 1.00 | 1.00 | 1.00 | 1.00 |
| SpCas9 | 1.00 | 1.80 | 1.02 | 1.04 | 0.85 | 0.96 | 1.05 | 0.94 | 0.96 | 0.97 | 0.89 | 1.02 | 1.20 | 0.94 | 0.98 | 1.15 | 0.98 | 1.08 | 0.99 | 1.05 | 1.00 | 1.01 | 1.05 | 1.01 | 1.05 | 0.99 |
| SaCas9 | 1.00 | 0.93 | 5.68 | 1.00 | 0.99 | 1.45 | 0.91 | 0.90 | 0.88 | 0.99 | 1.05 | 1.06 | 0.89 | 0.92 | 0.95 | 0.96 | 0.95 | 0.91 | 0.94 | 0.98 | 1.00 | 0.94 | 1.01 | 0.89 | 0.91 | 0.91 |
| CjCas9 | 1.00 | 0.89 | 0.97 | 5.61 | 1.06 | 1.01 | 0.91 | 0.90 | 1.49 | 0.96 | 1.10 | 0.96 | 0.86 | 0.89 | 0.94 | 1.00 | 0.96 | 0.93 | 1.06 | 1.03 | 1.01 | 0.98 | 0.97 | 0.91 | 0.96 | 1.01 |
| NmCas9 | 1.00 | 0.96 | 1.05 | 0.94 | 3.16 | 1.05 | 0.97 | 1.04 | 0.95 | 0.90 | 2.47 | 0.98 | 0.90 | 0.92 | 0.96 | 0.96 | 1.05 | 0.94 | 1.10 | 1.01 | 0.97 | 0.98 | 0.97 | 0.91 | 1.03 | 0.95 |
| St1Cas9 | 1.00 | 0.99 | 3.82 | 1.09 | 1.02 | 2.39 | 0.93 | 0.86 | 0.92 | 1.00 | 1.12 | 2.38 | 0.91 | 1.03 | 1.03 | 1.00 | 0.98 | 0.95 | 1.09 | 1.08 | 1.01 | 1.06 | 0.98 | 0.90 | 1.03 | 2.06 |
| FnCas9 | 1.00 | 0.92 | 1.04 | 1.07 | 1.13 | 0.99 | 3.61 | 0.93 | 0.88 | 0.99 | 1.33 | 0.96 | 0.89 | 0.97 | 1.00 | 1.02 | 1.08 | 0.95 | 1.05 | 1.03 | 1.01 | 1.00 | 0.96 | 0.95 | 0.97 | 0.96 |
| CdCas9 | 1.00 | 1.11 | 1.15 | 1.13 | 1.51 | 1.11 | 1.03 | 1.56 | 1.06 | 1.01 | 1.34 | 1.13 | 0.93 | 1.03 | 1.06 | 1.17 | 1.09 | 1.12 | 1.12 | 1.14 | 1.10 | 1.16 | 1.18 | 1.04 | 1.17 | 1.44 |
| ClCas9 | 1.00 | 1.01 | 1.05 | 3.06 | 1.22 | 1.07 | 0.99 | 1.05 | 2.16 | 0.99 | 1.10 | 1.00 | 0.96 | 1.02 | 0.99 | 1.17 | 1.12 | 0.93 | 0.98 | 1.06 | 1.01 | 1.01 | 0.92 | 0.95 | 1.04 | 0.92 |
| PlCas9 | 1.00 | 1.07 | 1.26 | 1.17 | 1.54 | 1.25 | 1.07 | 1.16 | 1.00 | 1.31 | 1.40 | 1.11 | 1.03 | 1.09 | 1.05 | 1.16 | 1.04 | 1.05 | 1.32 | 1.19 | 1.25 | 1.19 | 1.24 | 1.07 | 1.29 | 1.27 |
| NcCas9 | 1.00 | 0.96 | 1.05 | 1.05 | 3.03 | 1.03 | 0.96 | 0.92 | 0.91 | 1.02 | 3.30 | 0.97 | 0.85 | 0.93 | 0.98 | 0.97 | 0.96 | 0.95 | 1.16 | 1.03 | 0.99 | 0.94 | 0.93 | 0.94 | 0.99 | 0.92 |
| SpaCas9 | 1.00 | 0.97 | 3.22 | 1.10 | 1.15 | 3.48 | 0.99 | 0.96 | 0.97 | 1.02 | 1.38 | 4.25 | 0.86 | 1.00 | 1.03 | 1.04 | 1.10 | 0.97 | 1.04 | 1.08 | 1.11 | 1.00 | 0.95 | 1.04 | 0.99 | 1.20 |
| St3Cas9 | 1.00 | 1.09 | 0.94 | 0.96 | 0.58 | 0.83 | 0.95 | 0.81 | 0.94 | 0.93 | 0.63 | 0.93 | 1.08 | 0.89 | 0.92 | 0.90 | 0.87 | 0.96 | 0.87 | 0.93 | 0.89 | 0.89 | 0.88 | 0.91 | 0.83 | 0.81 |
| AsCas12a | 1.00 | 1.03 | 1.24 | 1.23 | 1.35 | 1.11 | 1.06 | 1.14 | 1.14 | 1.04 | 1.35 | 1.09 | 0.93 | 5.13 | 5.39 | 5.15 | 4.40 | 0.99 | 1.14 | 1.12 | 1.14 | 1.14 | 1.17 | 1.11 | 1.11 | 1.47 |
| FnCas12a | 1.00 | 1.01 | 1.14 | 1.08 | 1.24 | 1.01 | 1.00 | 1.08 | 1.02 | 0.99 | 1.20 | 1.00 | 0.87 | 2.77 | 6.01 | 2.20 | 1.70 | 0.98 | 1.05 | 1.10 | 1.04 | 1.02 | 1.06 | 0.97 | 1.05 | 0.92 |
| LbCas12a | 1.00 | 1.07 | 1.27 | 1.32 | 1.36 | 1.22 | 1.11 | 1.07 | 1.15 | 1.13 | 1.49 | 1.14 | 0.96 | 2.96 | 2.07 | 6.28 | 2.46 | 1.09 | 1.21 | 1.22 | 1.15 | 1.19 | 1.24 | 1.10 | 1.18 | 1.15 |
| MbCas12a | 1.00 | 0.96 | 1.21 | 1.17 | 1.24 | 1.22 | 1.06 | 0.97 | 1.08 | 1.08 | 1.42 | 1.06 | 0.93 | 4.16 | 4.44 | 5.43 | 4.47 | 0.96 | 1.12 | 1.16 | 1.14 | 1.12 | 1.24 | 0.92 | 1.03 | 1.23 |
| AaCas12b | 1.00 | 0.87 | 1.06 | 1.07 | 1.16 | 1.10 | 0.96 | 0.94 | 0.92 | 0.95 | 1.42 | 1.05 | 0.80 | 0.92 | 0.96 | 1.01 | 0.99 | 2.49 | 4.73 | 1.54 | 1.01 | 1.00 | 1.02 | 1.00 | 1.03 | 0.96 |
| AkCas12b | 1.00 | 0.98 | 1.01 | 1.06 | 1.08 | 1.04 | 0.98 | 0.94 | 0.96 | 1.00 | 1.15 | 0.86 | 0.93 | 0.94 | 0.98 | 1.01 | 0.98 | 2.44 | 4.36 | 1.26 | 1.04 | 1.00 | 1.10 | 0.94 | 0.88 | 0.90 |
| BvCas12b | 1.00 | 0.89 | 0.98 | 1.03 | 0.76 | 0.88 | 1.00 | 0.86 | 0.88 | 0.95 | 0.93 | 0.95 | 1.09 | 0.96 | 0.93 | 1.01 | 0.89 | 0.98 | 1.06 | 2.39 | 0.97 | 1.04 | 1.02 | 0.91 | 0.84 | 0.93 |
| PspCas13b | 1.00 | 0.91 | 0.98 | 1.03 | 0.94 | 0.97 | 1.08 | 0.98 | 0.93 | 1.01 | 1.18 | 1.12 | 0.97 | 1.05 | 1.04 | 1.11 | 1.06 | 0.99 | 1.10 | 1.05 | 0.79 | 1.05 | 1.18 | 0.99 | 1.00 | 0.99 |
| PguCas13b | 1.00 | 0.82 | 0.94 | 0.98 | 0.85 | 0.98 | 0.87 | 0.85 | 0.85 | 0.89 | 1.10 | 0.96 | 0.93 | 0.97 | 0.96 | 0.96 | 0.95 | 0.90 | 0.99 | 1.00 | 1.05 | 1.13 | 1.07 | 0.84 | 0.89 | 0.97 |
| RanCas13b | 1.00 | 0.83 | 0.88 | 0.90 | 0.79 | 0.82 | 0.88 | 0.96 | 0.83 | 0.88 | 0.98 | 0.99 | 0.89 | 0.88 | 0.89 | 0.91 | 0.89 | 0.87 | 0.91 | 0.99 | 1.01 | 1.91 | 2.01 | 0.83 | 0.91 | 0.89 |
| CasRx | 1.00 | 0.86 | 0.95 | 0.96 | 1.03 | 1.04 | 0.95 | 0.93 | 0.86 | 1.02 | 1.25 | 0.94 | 0.86 | 0.97 | 0.93 | 0.97 | 0.96 | 0.93 | 1.09 | 1.01 | 1.04 | 0.97 | 1.05 | 4.88 | 0.94 | 0.93 |
| PlmCasX | 1.00 | 0.91 | 0.95 | 1.05 | 0.97 | 1.03 | 0.93 | 0.96 | 0.89 | 1.00 | 1.22 | 0.99 | 0.87 | 0.94 | 0.98 | 0.99 | 1.00 | 0.93 | 1.11 | 1.09 | 1.08 | 1.06 | 1.08 | 0.96 | 4.16 | 0.95 |
| Cas14a1 | 1.00 | 0.94 | 1.06 | 1.16 | 0.92 | 0.96 | 1.18 | 0.90 | 0.91 | 1.01 | 1.00 | 0.96 | 0.88 | 0.95 | 1.04 | 1.06 | 0.96 | 0.94 | 1.05 | 1.10 | 1.16 | 1.07 | 1.15 | 0.96 | 0.91 | 1.83 |

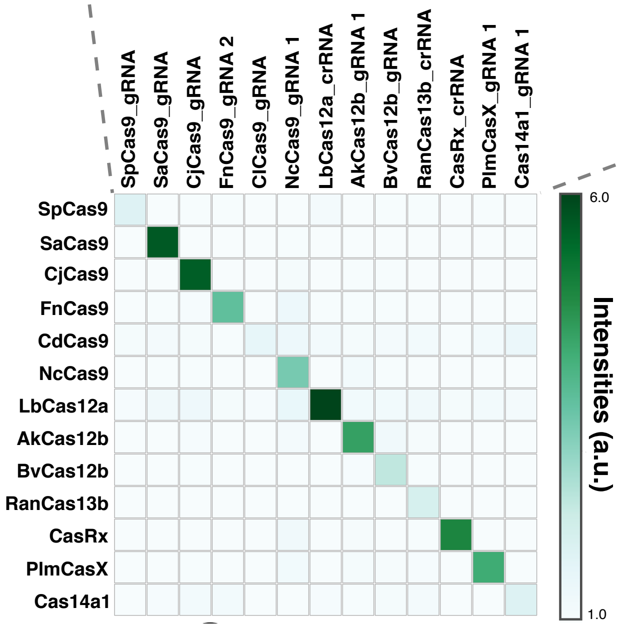

**Supplementary Figure 12|**

**Heat-map of 25 × 25 orthogonality matrix of representative Cas proteins and Cas-responsive ON switches.**

Values indicated the mean values in imaging analysis from three independent experiments performed on different days. Pruned images in the bottom indicate mutually orthogonal sets.

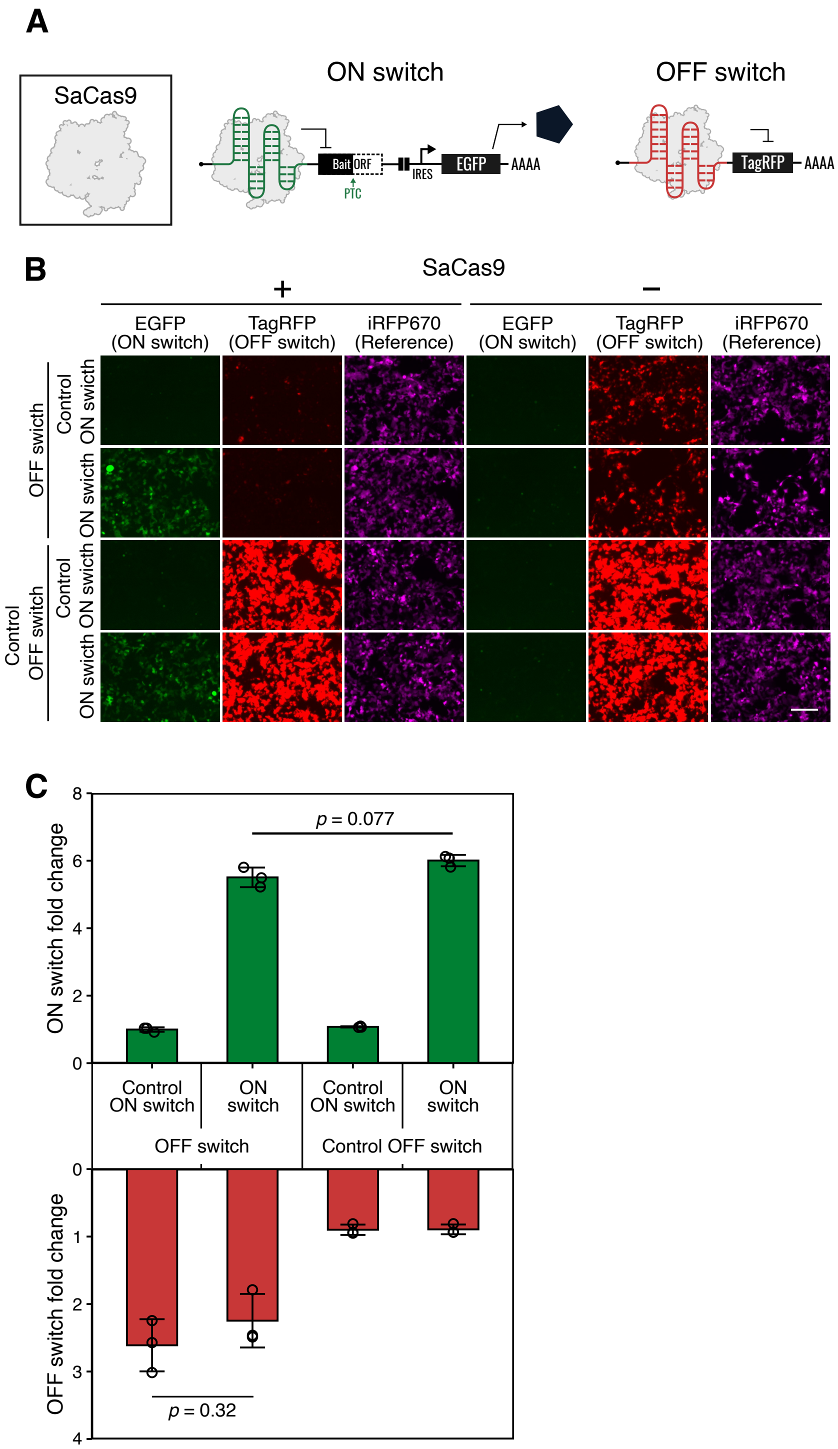
A
SaCas9
ON switch
OFF switch
Bait
ORF
PTC
IRES
EGFP
AAAA
TagRFP
AAAA
B
SaCas9
+
−
EGFP (ON switch)
TagRFP (OFF switch)
iRFP670 (Reference)
EGFP (ON switch)
TagRFP (OFF switch)
iRFP670 (Reference)
OFF swicth
Control ON swicth
ON swicth
Control OFF switch
Control ON swicth
ON swicth
C
ON switch fold change
p = 0.077
Control ON switch
ON switch
Control ON switch
ON switch
OFF switch
Control OFF switch
OFF switch fold change
p = 0.32

**Supplementary Figure 13|**

**Simultaneous regulation of translational activation and repression with SaCas9.**

(A) The schematic diagram of the simultaneous translational regulation system. SaCas9 was used as the trigger protein. Translational activation and repression were monitored with EGFP and TagRFP, respectively. (B) Fluorescent microscopic images in each condition. Translational activation and repression were observed simultaneously when SaCas9 was transfected. Scale bar, 200 μm. (C) Quantitative data of reporter expression levels. The data were obtained by imaging analysis. iRFP670 was co-transfected as a reference and used to define a transfection-positive area. OFF switch: a plasmid for expressing TagRFP whose translation is repressed by SaCas9. ON switch: a plasmid for expressing EGFP whose translation is activated by SaCas9. Control OFF and ON switch: plasmids without the gRNA sequence for SaCas9. Error bars represent mean ± SD from three independent experiments. Statistical analyses were carried out with unpaired two-tailed Student's t-test.

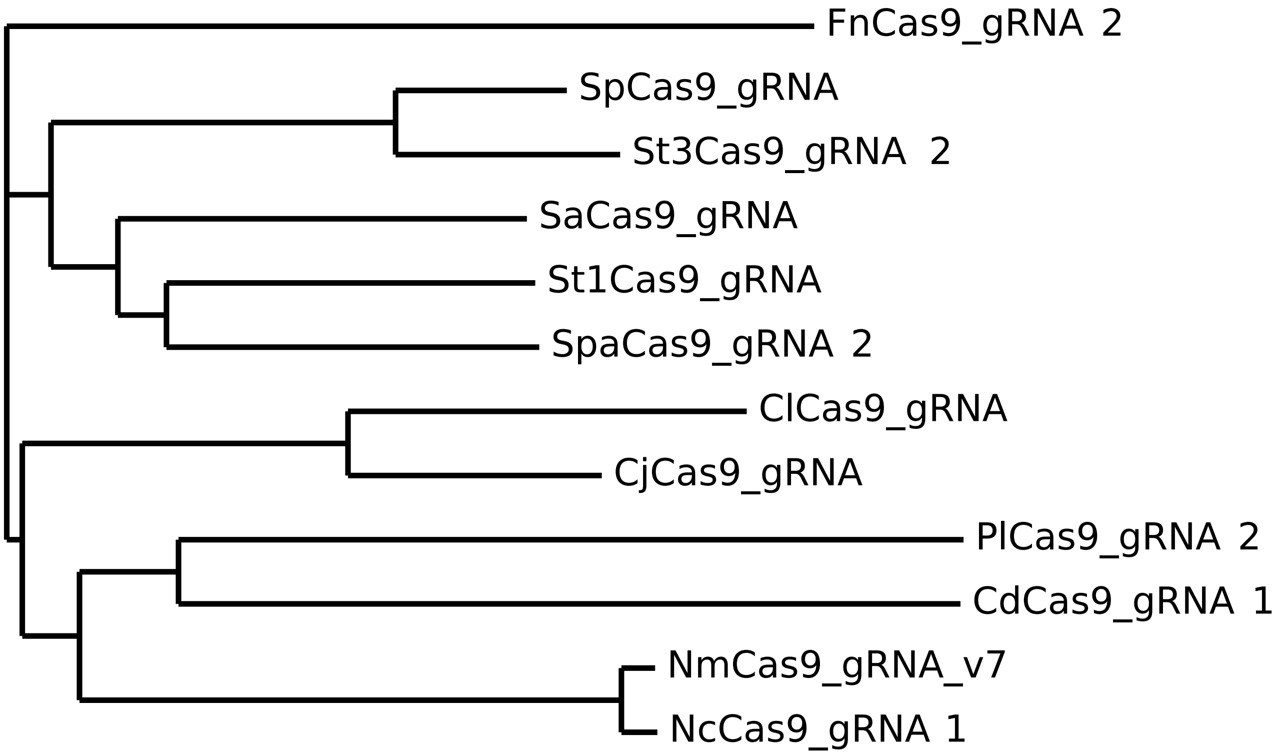

**Supplementary Figure 14|**

**Phylogenetic tree of Cas9 gRNA used in OFF switch orthogonality.**

The sequences were aligned with the MUSCLE web tool. The pairs that showed crosstalk were grouped in closer clusters.

**Supplementary Table 1**

Information of Cas proteins used in this study.

| | Name | Origin | RNA-guided nuclease activity in mammalian cells | Repression efficiency (%) | Size (bp)*1 | gRNA length (nt)*2 |
|---|---|---|---|---|---|---|
| 1 | SpCas9 | *Streptococcus pyogenes* | + | 85 | 4140 | 97 |
| 2 | SaCas9 | *Staphylococcus aureus* | + | 93 | 3345 | 97 |
| 3 | CjCas9 | *Campylobacter jejuni* | + | 81 | 3012 | 93 |
| 4 | NmCas9 | *Neisseria meningitidis* | + | v0: 1, v1: 5, v2: 0, v3: 58, v4: 57, v5: 28, v6: 45, v7: 62 | 3333 | 121 |
| 5 | St1Cas9 | *Streptococcus thermophilus* | + | 81 | 3402 | 147 |
| 6 | FnCas9 | *Francisella novicida* | + (with proxy-CRISPR) | 1: 41, 2: 90 | 4950 | 1: 75, 2: 111 |
| 7 | CdCas9 | *Corynebacterium diphtheriae* | + | 1:26, 2:19 | 3318 | 1: 98, 2: 97 |
| 8 | ClCas9 | *Campylobacter lari* CF89-12 | - | 36 | 3075 | 113 |
| 9 | PlCas9 | *Parvibaculum lavamentivorans* | - | 1: 13, 2: 29 | 3177 | 1: 122, 2: 124 |
| 10 | NcCas9 | *Neisseria cinerea* | + (with proxy-CRISPR) | 1: 88, 2: 83 | 3312 | 1: 108, 2: 162 |
| 11 | SpaCas9 | *Streptococcus pasteurianus* | + | 1: 77, 2: 84 | 3456 | 1: 83, 2: 82 |
| 12 | St3Cas9 | *Streptococcus thermophilus* | + | 1: 87, 2: 87 | 4260 | 1: 80, 2: 94 |

| | Name | Origin | RNA-guided nuclease activity in mammalian cells | Repression efficiency (%) | Size (bp)*1 | gRNA length (nt)*2 |
|---|---|---|---|---|---|---|
| 1 | AsCpf1 (AsCas12a) | *Acidaminococcus sp.* BV3L6 | + | 90 | 4062 | 43 |
| 2 | FnCpf1 (FnCas12a) | *Francisella novicida* U112 | + | 86 | 4038 | 20 |
| 3 | LbCpf1 (LbCas12a) | *Lachnospiraceae bacterium* ND2006 | + | 91 | 3822 | 21 |
| 4 | MbCpf1 (MbCas12a) | *Moraxella bovoculi* 237 | + | 93 | 4257 | 21 |
| 5 | AaCas12b | *Alicyclobacillus acidiphilus* | + | 83 | 3447 | 137 |
| 6 | AkCas12b | *Alicyclobacillus kakegawensis* | + | 1: 91, 2: 78 | 3597 | 1: 117, 2: 137 |
| 7 | BvCas12b | *Bacillus sp.* NSP2.1 | + | 85 | 3522 | 98 |

| | Name | Origin | RNA-guided nuclease activity in mammalian cells | Repression efficiency (%) | Size (bp)*1 | gRNA length (nt)*2 |
|---|---|---|---|---|---|---|
| 1 | PspCas13b | *Prevotella sp.* | + | 97 | 3429 | 66 |
| 2 | PguCas13b | *Porphyromonas gulae* | + | 96 | 3657 | 66 |
| 3 | RanCas13b | *Riemerella anatipestifer* | + | 83 | 3417 | 66 |
| 4 | CasRx (RfxCas13d) | *Ruminococcus flavefaciens XPD3002* | + | 87 | 3003 | 58 |

---

| | Name | Origin | RNA-guided nuclease activity in mammalian cells | Repression efficiency (%) | Size (bp)*1 | gRNA length (nt)*2 |
|---|---|---|---|---|---|---|
| 1 | PlmCasX | *Planctomycetes* | + | 1: 63, 2: 66 | 2937 | 1: 122, 2: 102 |
| 2 | Cas14a1 | Uncultured archaeron | + | 1: 86, 2: 78 | 1590 | 1: 163, 2: 204 |

*1 The length of the ORF used in this study (including stop codon)

*2 The length of the gRNA/crRNA used in this study

**Supplementary Table 2**

Profile of the 60 AND gates in Figure 5.

| AND # | Input A | Input B | Cas protein C |
|---|---|---|---|
| 1 | PguCas13b | MbCas12a | PspCas13b |
| 2 | PguCas13b | SaCas9 | PspCas13b |
| 3 | PguCas13b | AkCas12b | PspCas13b |
| 4 | PguCas13b | NcCas9 | PspCas13b |
| 5 | MbCas12a | SaCas9 | PspCas13b |
| 6 | MbCas12a | AkCas12b | PspCas13b |
| 7 | MbCas12a | NcCas9 | PspCas13b |
| 8 | SaCas9 | AkCas12b | PspCas13b |
| 9 | SaCas9 | NcCas9 | PspCas13b |
| 10 | AkCas12b | NcCas9 | PguCas13b |
| 11 | PspCas13b | MbCas12a | PguCas13b |
| 12 | PspCas13b | SaCas9 | PguCas13b |
| 13 | PspCas13b | AkCas12b | PguCas13b |
| 14 | PspCas13b | NcCas9 | PguCas13b |
| 15 | MbCas12a | SaCas9 | PguCas13b |
| 16 | MbCas12a | AkCas12b | PguCas13b |
| 17 | MbCas12a | NcCas9 | PguCas13b |
| 18 | SaCas9 | AkCas12b | PguCas13b |
| 19 | SaCas9 | NcCas9 | PguCas13b |
| 20 | AkCas12b | NcCas9 | PguCas13b |
| 21 | PspCas13b | PguCas13b | MbCas12a |
| 22 | PspCas13b | SaCas9 | MbCas12a |
| 23 | PspCas13b | AkCas12b | MbCas12a |
| 24 | PspCas13b | NcCas9 | MbCas12a |
| 25 | PguCas13b | SaCas9 | MbCas12a |
| 26 | PguCas13b | AkCas12b | MbCas12a |
| 27 | PguCas13b | NcCas9 | MbCas12a |
| 28 | SaCas9 | AkCas12b | MbCas12a |
| 29 | SaCas9 | NcCas9 | MbCas12a |
| 30 | AkCas12b | NcCas9 | MbCas12a |

| AND # | Input A | Input B | Cas protein C |
|---|---|---|---|
| 31 | PspCas13b | PguCas13b | SaCas9 |
| 32 | PspCas13b | MbCas12a | SaCas9 |
| 33 | PspCas13b | AkCas12b | SaCas9 |
| 34 | PspCas13b | NcCas9 | SaCas9 |
| 35 | PguCas13b | MbCas12a | SaCas9 |
| 36 | PguCas13b | AkCas12b | SaCas9 |
| 37 | PguCas13b | NcCas9 | SaCas9 |
| 38 | MbCas12a | AkCas12b | SaCas9 |
| 39 | MbCas12a | NcCas9 | SaCas9 |
| 40 | AkCas12b | NcCas9 | SaCas9 |
| 41 | PspCas13b | PguCas13b | AkCas12b |
| 42 | PspCas13b | MbCas12a | AkCas12b |
| 43 | PspCas13b | SaCas9 | AkCas12b |
| 44 | PspCas13b | NcCas9 | AkCas12b |
| 45 | PguCas13b | MbCas12a | AkCas12b |
| 46 | PguCas13b | SaCas9 | AkCas12b |
| 47 | PguCas13b | NcCas9 | AkCas12b |
| 48 | MbCas12a | SaCas9 | AkCas12b |
| 49 | MbCas12a | NcCas9 | AkCas12b |
| 50 | SaCas9 | NcCas9 | AkCas12b |
| 51 | PspCas13b | PguCas13b | NcCas9 |
| 52 | PspCas13b | MbCas12a | NcCas9 |
| 53 | PspCas13b | SaCas9 | NcCas9 |
| 54 | PspCas13b | AkCas12b | NcCas9 |
| 55 | PguCas13b | MbCas12a | NcCas9 |
| 56 | PguCas13b | SaCas9 | NcCas9 |
| 57 | PguCas13b | AkCas12b | NcCas9 |
| 58 | MbCas12a | SaCas9 | NcCas9 |
| 59 | MbCas12a | AkCas12b | NcCas9 |
| 60 | SaCas9 | AkCas12b | NcCas9 |

**Supplementary Table 3**

Key plasmids used in this study.

| Description | Original name | Benchling link | Short name | Figure |
|---|---|---|---|---|
| OFF switch | pGluc-Sp_gRNA-EGFP | https://benchling.com/s/seq-YB5Dr4uWTfanMOKvIUsY | Sp_gRNA or SpCas9_gRNA | Fig1, 2, 3, FigS2, S3, S9, S10 |
| OFF switch | pGluc-Sa_gRNA-EGFP | https://benchling.com/s/seq-M1VrldwdMbyqmlAdm2TA | Sa_gRNA or SaCas9_gRNA | Fig1, 2, 3, 5, FigS2, S10 |
| OFF switch | pGluc-Cj_gRNA-EGFP | https://benchling.com/s/seq-i683ARLbCYEjQdq5Cnh0 | CjCas9_gRNA | Fig1, 3, FigS2, S10 |
| OFF switch | pGluc-St1_gRNA-EGFP | https://benchling.com/s/seq-TskVXFHofI9E2pxSQWkM | St1Cas9_gRNA | Fig1, 3, FigS2, S10 |
| OFF switch | pFn_gRNA_v1-EGFP | https://benchling.com/s/seq-ovf2dJM05eLri2RCfAHh | FnCas9_gRNA 1 | Fig1, FigS2 |
| OFF switch | pFn_gRNA_v2-EGFP | https://benchling.com/s/seq-m3xNCSEq42TLjiLL293S | FnCas9_gRNA 2 | Fig1, 3, FigS2, S10 |
| OFF switch | pCd_gRNA_v1-EGFP | https://benchling.com/s/seq-K6uhc21sAPxxOEK6PZuw | CdCas9_gRNA 1 | Fig1, 3, FigS2, S10 |
| OFF switch | pCd_gRNA_v2-EGFP | https://benchling.com/s/seq-KZVPW44Dk4V6u4zafopV | CdCas9_gRNA 2 | Fig1, FigS2 |
| OFF switch | pCl_gRNA-EGFP | https://benchling.com/s/seq-H4UxWLOhgNvA0YR4Szip | ClCas9_gRNA | Fig1, 3, FigS2, S10 |
| OFF switch | pPl_gRNA_v1-EGFP | https://benchling.com/s/seq-VfDteXn2JGE7dYPfy9D3 | PlCas9_gRNA 1 | Fig1, FigS2 |
| OFF switch | pPl_gRNA_v2-EGFP | https://benchling.com/s/seq-wilfLbHEW4cECDThdcwt | PlCas9_gRNA 2 | Fig1, 3, FigS2, S10 |
| OFF switch | pNc_gRNA_v1-EGFP | https://benchling.com/s/seq-9GsfCiNp0dlL6B6UFmpP | NcCas9_gRNA 1 | Fig1, 3, 5, FigS2 |
| OFF switch | pNc_gRNA_v2-EGFP | https://benchling.com/s/seq-oF6McTKt84TdwsJ5Wt1H | NcCas9_gRNA 2 | Fig1, FigS2 |
| OFF switch | pSpa_gRNA_v1-EGFP | https://benchling.com/s/seq-PRk8eHaoV3JHCtXrX3BE | SpaCas9_gRNA 1 | Fig1, FigS2 |
| OFF switch | pSpa_gRNA_v2-EGFP | https://benchling.com/s/seq-Oh0lQ13b1xRVbmpRBqYH | SpaCas9_gRNA 2 | Fig1, 3, FigS2, S10 |
| OFF switch | pGluc-St3_gRNA_v1-EGFP | https://benchling.com/s/seq-N17o8zbHO751wm0LPAsa | St3Cas9_gRNA 1 | Fig1, FigS2 |
| OFF switch | pGluc-St3_gRNA_v2-EGFP | https://benchling.com/s/seq-BwCIYQ2YCuw2O3GdfvUV | St3Cas9_gRNA 2 | Fig1, 3, FigS2, S10 |
| OFF switch | pGluc-AsCas12a_crRNA-EGFP | https://benchling.com/s/seq-glNfU5jZtiKihoYUIYbg | AsCas12a_crRNA | Fig1, 3, FigS2, S6, S10 |
| OFF switch | pFnCas12a_crRNA-EGFP | https://benchling.com/s/seq-V8aaKQn6IXZBvwOrjF4e | FnCas12a_crRNA | Fig1, 3, FigS2, S10 |
| OFF switch | pLbCas12a_crRNA-EGFP | https://benchling.com/s/seq-AjWByTr164WVX6iEujJH | LbCas12a_crRNA | Fig1, 3, FigS2, S10 |
| OFF switch | pMbCas12a_crRNA-EGFP | https://benchling.com/s/seq-hbYhHvSP2IG6F4WpK4IN | MbCas12a_crRNA | Fig1, 3, 5, FigS2, S10 |
| OFF switch | pAaCas12b_sgRNA-EGFP | https://benchling.com/s/seq-0ah7ki0EgntdfZahBDPU | AaCas12a_crRNA | Fig1, 3, FigS2, S10 |
| OFF switch | pAkCas12b_sgRNA_v1-EGFP | https://benchling.com/s/seq-7ZiHjayLdyOwjiBwW9qe | AkCas12b_gRNA 1 | Fig1, 3, 5, FigS2, S10 |
| OFF switch | pAkCas12b_sgRNA_v2-EGFP | https://benchling.com/s/seq-WhxCmn2hJG2pYh5T4JaK | AkCas12b_gRNA 2 | Fig1, FigS2 |
| OFF switch | pBvCas12b_sgRNA-EGFP | https://benchling.com/s/seq-ERy3edQQ4oRn27cBoE1F | BvCas12b_gRNA | Fig1, 3, FigS2, S10 |
| OFF switch | pGluc-Psp_crRNA-EGFP | https://benchling.com/s/seq-WJlEdGEGMXhOOPGzuGHZ | PspCas13b_crRNA | Fig1, 3, 5, FigS2, S10 |
| OFF switch | pGluc-Pgu_crRNA-EGFP | https://benchling.com/s/seq-q7r7cKeYiylzuEg4k7bP | PguCas13b_crRNA | Fig1, 3, 5, FigS2, S10 |
| OFF switch | pGluc-Ran_crRNA-EGFP | https://benchling.com/s/seq-dR5ZevWPdwURmwnkXMsA | RanCas13b_crRNA | Fig1, 3, FigS2, S10 |
| OFF switch | pGluc-CasRX_crRNA-EGFP | https://benchling.com/s/seq-E6RZI3LyVrpYHNZuhz06 | CasRx_crRNA | Fig1, 3, FigS2, S10 |

| | | | | |
|---|---|---|---|---|
| OFF switch | pPlmCasX_gRNA-EGFP | https://benchling.com/s/seq-CN4z7nmpzAYbyQPomM4H | PlmCasX_gRNA 1 | Fig1, FigS2 |
| OFF switch | pPlmCasX_gRNA(dSpacer)-EGFP | https://benchling.com/s/seq-XSQeSp220dxaYoc5BkL9 | PlmCasX_gRNA 2 | Fig1, 3, FigS2, S10 |
| OFF switch | pCas14a1_sgRNA1-EGFP | https://benchling.com/s/seq-nL3klfKUsOsPT2W8iuz7 | Cas14a1_gRNA 1 | Fig1, 3, FigS2, S10 |
| OFF switch | pCas14a1_sgRNA2-EGFP | https://benchling.com/s/seq-oIcaD1LAx1CS6wGriPSc | Cas14a1_gRNA 2 | Fig1, FigS2 |
| OFF switch | pGluc-Nm_gRNA-EGFP | https://benchling.com/s/seq-9ShtAQING48KU8WRPZI0 | NmCas9_gRNA_v0 | Fig1, FigS2, S5 |
| OFF switch | pGluc-Nm_gRNA-STOP-EGFP | https://benchling.com/s/seq-I3yoBcMcdBdqRQRUwmXg | NmCas9_gRNA_v1 | Fig1, FigS2, S5 |
| OFF switch | pGluc-Nm_gRNA_v2-EGFP | https://benchling.com/s/seq-4QVuutAH6V2urfJllMRy | NmCas9_gRNA_v2 | Fig1, FigS2, S5 |
| OFF switch | pGluc-Nm_gRNA_v3-EGFP | https://benchling.com/s/seq-wbDRU1SvlenI79ZS1H2F | NmCas9_gRNA_v3 | Fig1, FigS2, S5 |
| OFF switch | pNm_gRNA_v4-EGFP | https://benchling.com/s/seq-b1iBbNY6afS8lW4wlZMq | NmCas9_gRNA_v4 | Fig1, FigS2, S5 |
| OFF switch | pNm_gRNA_v5-EGFP | https://benchling.com/s/seq-mPrZo0zK3tP8fOo0fXtN | NmCas9_gRNA_v5 | Fig1, FigS2, S5 |
| OFF switch | pNm_gRNA_v6-EGFP | https://benchling.com/s/seq-tHODiWmkQAncUaHAP2Et | NmCas9_gRNA_v6 | Fig1, FigS2, S5 |
| OFF switch | pNm_gRNA_v7-EGFP | https://benchling.com/s/seq-CXq80mUzT3gHGyDvJXiX | NmCas9_gRNA_v7 | Fig1, 3, FigS2, S5, S10 |
| Trigger | pcDNA3.1-SpCas9 | https://benchling.com/s/seq-EmEEkWHXs3Jf4rcdpaTg | SpCas9/WT | Fig1, 2, 3, Fig S2, S3, S7, S9, S10 |
| Trigger | pcDNA3.1+-SaCas9 | https://benchling.com/s/seq-Sj4R81pj6RXIZ6Jly9p4 | SaCas9 | Fig1, 2, 3, 5, Fig S2, S7, S10, S13 |
| Trigger | pcDNA3.1+-CjCas9 | https://benchling.com/s/seq-6eXuV9JcFQeQbmYwfJX8 | CjCas9 | Fig1, 3, Fig S2, S7, S10 |
| Trigger | pcDNA3.1+-NmCas9 | https://benchling.com/s/seq-Wfg75vjX32YQQJkiyuYE | NmCas9 | Fig1, 3, Fig S2, S5, S7, S10 |
| Trigger | pcDNA3.1+-st1Cas9 | https://benchling.com/s/seq-VRWF7oQowjQbAvG8mYce | St1Cas9 | Fig1, 3, Fig S2, S7, S10 |
| Trigger | pcDNA3.1+-FnCas9 | https://benchling.com/s/seq-T1yYZahdAytxkLbQf5Jw | FnCas9 | Fig1, 3, Fig S2, S7, S10 |
| Trigger | pcDNA3.1+-CdCas9 | https://benchling.com/s/seq-kC3slahy2rNyCbP39Uls | CdCas9 | Fig1, 3, Fig S2, S7, S10 |
| Trigger | pcDNA3.1+-ClCas9 | https://benchling.com/s/seq-QMBrgPUM4wQ6nJFQp2Jv | ClCas9 | Fig1, 3, Fig S2, S7, S10 |
| Trigger | pcDNA3.1+-PlCas9 | https://benchling.com/s/seq-ibuHG9biYiK2zC93XPUq | PlCas9 | Fig1, 3, Fig S2, S7, S10 |
| Trigger | pcDNA3.1+-NcCas9 | https://benchling.com/s/seq-ifoU1HSjWp7QaJ2rnz4O | NcCas9 | Fig1, 3, 5, Fig S2, S7, S10 |
| Trigger | pcDNA3.1+-SpaCas9 | https://benchling.com/s/seq-hmk13W570AKm7GQ0roes | SpaCas9 | Fig1, 3, Fig S2, S7, S10 |
| Trigger | pcDNA3.1+-St3Cas9 | https://benchling.com/s/seq-2q6Vcj3AwAHrO3ZXoP3Y | St3Cas9 | Fig1, 3, Fig S2, S7, S10 |
| Trigger | pcDNA3.1+-AsCpf1 | https://benchling.com/s/seq-7cpBK5lcPWKG0wk88fqZ | AsCas12a | Fig1, 3, Fig S2, S6, S7, S10 |
| Trigger | pcDNA3.1-hMbCpf1 | https://benchling.com/s/seq-Ap2WTz89rTXvcdSrQkgT | MbCas12a | Fig1, 3, 5, Fig S2, S7, S10 |
| Trigger | pcDNA3.1+-AaCas12b | https://benchling.com/s/seq-Cg0Aurq89nwhVyoOf7cz | AaCas12b | Fig1, 3, Fig S2, S7, S10 |
| Trigger | pcDNA3.1-hFnCpf1 | https://benchling.com/s/seq-NQNXT0q4aU2jFTtNf3Oq | FnCas12a | Fig1, 3, Fig S2, S7, S10 |
| Trigger | pcDNA3.1-hLbCpf1 | https://benchling.com/s/seq-LRXskpS65qeRNfMcOMZT | LbCas12a | Fig1, 3, Fig S2, S7, S10 |
| Trigger | pcDNA3.1+-AkCas12b | https://benchling.com/s/seq-9CJWprWiG5TviPN6Zh4R | AkCas12b | Fig1, 3, 5, Fig S2, S7, S10 |

| | | | | |
|---|---|---|---|---|
| Trigger | pcDNA3.1+-BvCas12b | https://benchling.com/s/seq-lwRpPTKuL6fB3m0XNzK6 | BvCas12b | Fig1, 3, Fig S2, S7, S10 |
| Trigger | pcDNA3.1-PspCas13b(WT)-NES-myc-His6 | https://benchling.com/s/seq-X0bNQN8K5SsJBN6F3OOX | PspCas13b | Fig1, 3, 5, Fig S2, S7, S10 |
| Trigger | pcDNA3.1+-PguCas13b-NES | https://benchling.com/s/seq-BPSiSwCOpWBOxS0GGgcD | PguCas13b | Fig1, 3, 5, Fig S2, S7, S10 |
| Trigger | pcDNA3.1+-RanCas13b-NES | https://benchling.com/s/seq-j8EjvpKgsQntC2aumCLE | RanCas13b | Fig1, 3, Fig S2, S7, S10 |
| Trigger | pcDNA3.1+-CasRx | https://benchling.com/s/seq-9sXxgMXbXXXfw9qoAtLo | CasRx | Fig1, 3, Fig S2, S7, S10 |
| Trigger | pcDNA3.1+-PlmCasX | https://benchling.com/s/seq-iwnSWt8Fk44t9U9QBnqa | PlmCasX | Fig1, 3, Fig S2, S7, S10 |
| Trigger | pcDNA3.1+-Cas14a1 | https://benchling.com/s/seq-omwxft6FEDKBWlKxejS4 | Cas14a1 | Fig1, 3, Fig S2, S7, S10 |
| Trigger | pcDNA3.1-SpCas9(D10A) | https://benchling.com/s/seq-rglwwC4Z2fEgTNhxNCmX | D10A | Fig S3 |
| Trigger | pcDNA3.1-SpCas9(D10A_H840) | https://benchling.com/s/seq-oTpkJKDiTUBXWxa45FDX | D10A, H840A | Fig S3 |
| Trigger | pcDNA3.1-SpCas9(dNLS) | https://benchling.com/s/seq-xIMuZsE3gLNDmcQDstVq | ΔNLS | Fig S3 |
| Acr | pcDNA3.1+-AcrIIA4 | https://benchling.com/s/seq-gGe8r2DZoDQvxqDhar3j | AcrIIA4 | Fig S3 |
| Trigger | pcDNA3.1+-AsCpf1(H800A) | https://benchling.com/s/seq-9U2OSQeiLx0biaTx8INe | AsCas12a(H800A) | Fig S6 |
| ON switch | pNMD-ON-Gluc-Sp_gRNA-EGFP | https://benchling.com/s/seq-rBiY9xY8IxCQlmCDzb2v | SpCas9_gRNA | Fig 1, Fig S7, S11, S12, S13 |
| ON switch | pNMD-ON-Gluc-Sa_gRNA | https://benchling.com/s/seq-Rkh4phoiHfg2TfGdl4LP | SaCas9_gRNA | Fig 1, Fig S7, S11, S12, S13 |
| ON switch | pNMD-ON-Gluc-Cj_gRNA | https://benchling.com/s/seq-ytcIQFC5hKdeTzpgVAgg | CjCas9_gRNA | Fig 1, Fig S7, S11, S12 |
| ON switch | pNMD-ON-Gluc-St1_gRNA | https://benchling.com/s/seq-edpM2PVwBHJPhFaMwMGq | St1Cas9_gRNA | Fig 1, Fig S7, S11, S12 |
| ON switch | pNMD-ON-Gluc_Nm_gRNA_v1 | https://benchling.com/s/seq-j8yTztzbcE5l6BnnaYcV | NmCas9_gRNA 1 | Fig S7 |
| ON switch | pNMD-ON-Gluc_Nm_gRNA_v2 | https://benchling.com/s/seq-4JkdlEgacfZVliRF9dFW | NmCas9_gRNA 2 | Fig S7 |
| ON switch | pNMD-ON-Gluc_Nm_gRNA_v3 | https://benchling.com/s/seq-AYCFLt1uYmg1tnfU7TLk | NmCas9_gRNA 3 | Fig 1, Fig S7, S11, S12 |
| ON switch | pNMD-ON-Gluc_Nm_gRNA_v4 | https://benchling.com/s/seq-lg2jkJ3G8H1fd4MMQ4CO | NmCas9_gRNA 4 | Fig S7 |
| ON switch | pNMD-ON-Gluc_Nm_gRNA_v5 | https://benchling.com/s/seq-fXdYnMeOXbnz6Whw4Jed | NmCas9_gRNA 5 | Fig S7 |
| ON switch | pNMD-ON-Gluc_Nm_gRNA_v6 | https://benchling.com/s/seq-VX81PClbzWVwRpJdbQNJ | NmCas9_gRNA 6 | Fig S7 |
| ON switch | pNMD-ON-Gluc_Nm_gRNA_v7 | https://benchling.com/s/seq-gzSPC4IFm9ll9ShoyHIr | NmCas9_gRNA 7 | Fig S7 |
| ON switch | pNMD-ON-Fn_gRNA_v1 | https://benchling.com/s/seq-UpTK1Av0CTE5Qzgf5OSR | FnCas9_gRNA 1 | Fig S7 |
| ON switch | pNMD-ON-Fn_gRNA_v2 | https://benchling.com/s/seq-wnexJ24lCiNDjWyIaPjl | FnCas9_gRNA 2 | Fig 1, Fig S7, S11, S12 |
| ON switch | pNMD-ON-Cd_gRNA_v1 | https://benchling.com/s/seq-aZ03Yf3zGnKi9y1qrP35 | CdCas9_gRNA 1 | Fig S7 |
| ON switch | pNMD-ON-Cd_gRNA_v2 | https://benchling.com/s/seq-Mssi3187k0LiwSpbSY1g | CdCas9_gRNA 2 | Fig 1, Fig S7, S11, S12 |
| ON switch | pNMD-ON-Cl_gRNA | https://benchling.com/s/seq-gkRuH0CD6l8VFkLfGTuN | ClCas9_gRNA | Fig 1, Fig S7, S11, S12 |
| ON switch | pNMD-ON-Pl_gRNA_v1 | https://benchling.com/s/seq-neQ5XOaKZayZxaEnBpBJ | PlCas9_gRNA 1 | Fig S7 |

| | | | | |
|---|---|---|---|---|
| ON switch | pNMD-ON-Pl_gRNA_v2 | https://benchling.com/s/seq-uo12MN8p9ELYe4KBwKPy | PlCas9_gRNA 2 | Fig 1, Fig S7, S11, S12 |
| ON switch | pNMD-ON-Nc_gRNA_v1 | https://benchling.com/s/seq-57djv04vQeZYOqLdpZB9 | NcCas9_gRNA 1 | Fig 1, Fig S7, S11, S12 |
| ON switch | pNMD-ON-Nc_gRNA_v2 | https://benchling.com/s/seq-5Kd4BkOffc6EGynJb7Xf | NcCas9_gRNA 2 | Fig S7 |
| ON switch | pNMD-ON-Spa_gRNA_v1 | https://benchling.com/s/seq-fnoQ3X1Qs6sCtgmAhZQ6 | SpaCas9_gRNA 1 | Fig S7 |
| ON switch | pNMD-ON-Spa_gRNA_v2 | https://benchling.com/s/seq-OmmTbhPsOewdpW8J7vJh | SpaCas9_gRNA 2 | Fig 1, Fig S7, S11, S12 |
| ON switch | pNMD-ON-St3_gRNA_v1 | https://benchling.com/s/seq-vvJqP4bzkFr0jzUDum9m | St3Cas9_gRNA 1 | Fig 1, Fig S7, S11, S12 |
| ON switch | pNMD-ON-St3_gRNA_v2 | https://benchling.com/s/seq-0hRIB3bEUrvRH6pStcVP | St3Cas9_gRNA 2 | Fig S7 |
| ON switch | pNMD-ON-Gluc-AsCas12a_crRNA | https://benchling.com/s/seq-uMWXFaSnWI4Tj569dVR3 | AsCas12a_crRNA | Fig 1, Fig S7, S11, S12 |
| ON switch | pNMD-ON-FnCas12a_crRNA | https://benchling.com/s/seq-zCLlzSzy9FGZVE8LqSGw | FnCas12a_crRNA | Fig 1, Fig S7, S11, S12 |
| ON switch | pNMD-ON-LbCas12a_crRNA | https://benchling.com/s/seq-097cK15zRAFJnNdSzJrU | LbCas12a_crRNA | Fig 1, Fig S7, S11, S12 |
| ON switch | pNMD-ON-MbCas12a_crRNA | https://benchling.com/s/seq-2MgLIE9wFI3d5ljtMnbC | MbCas12a_crRNA | Fig 1, Fig S7, S11, S12 |
| ON switch | pNMD-ON-AaCas12b_sgRNA | https://benchling.com/s/seq-omJehIz6xnx7yEdD9GMw | AaCas12a_crRNA | Fig 1, Fig S7, S11, S12 |
| ON switch | pNMD-ON-AkCas12b_sgRNA_v1 | https://benchling.com/s/seq-eNnHdLLBxiWLCq3zFjEl | AkCas12b_gRNA 1 | Fig 1, Fig S7, S11, S12 |
| ON switch | pNMD-ON-AkCas12b_sgRNA_v2 | https://benchling.com/s/seq-9AOIST8qZ7TUMkhdqs9K | AkCas12b_gRNA 2 | Fig S7 |
| ON switch | pNMD-ON-BvCas12b_sgRNA | https://benchling.com/s/seq-y5azslDwpaymVWAunhno | BvCas12b_gRNA | Fig 1, Fig S7, S11, S12 |
| ON switch | pNMD-ON-Gluc-Psp_crRNA-EGFP | https://benchling.com/s/seq-6A2RLRkhHstAACjndh3T | PspCas13b_crRNA | Fig 1, Fig S7, S11, S12 |
| ON switch | pNMD-ON-Gluc-Pgu_crRNA | https://benchling.com/s/seq-A8zYUC0tnz18biHZwwih | PguCas13b_crRNA | Fig 1, Fig S7, S11, S12 |
| ON switch | pNMD-ON-Gluc_Ran_crRNA | https://benchling.com/s/seq-INRGbAowneL4z9J9NlpK | RanCas13b_crRNA | Fig 1, Fig S7, S11, S12 |
| ON switch | pNMD-ON-Gluc-CasRx_crRNA | https://benchling.com/s/seq-bHrydKbzUXtMT2df0vvF | CasRx_crRNA | Fig 1, Fig S7, S11, S12 |
| ON switch | pNMD-ON-PlmCasX_gRNA | https://benchling.com/s/seq-1ASZJgt3yMozw41vGMC4 | PlmCasX_gRNA 1 | Fig 1, Fig S7, S11, S12 |
| ON switch | pNMD-ON-PlmCasX_gRNA(dSpacer) | https://benchling.com/s/seq-2vhdu4FdsBUepnpg8sbH | PlmCasX_gRNA 2 | Fig S7 |
| ON switch | pNMD-ON-Cas14a1_sgRNA_v1 | https://benchling.com/s/seq-X7M8REytCChUfrThKnd0 | Cas14a1_gRNA 1 | Fig 1, Fig S7, S11, S12 |
| ON switch | pNMD-ON-Cas14a1_sgRNA_v2 | https://benchling.com/s/seq-gcwfVBOq93RuFqMvzOZm | Cas14a1_gRNA 2 | Fig S7 |
| Trigger | pcDNA3.1-SpCas9(1-713) | https://benchling.com/s/seq-WhnSX12uuWktk1Ptnl6N | SpCas9(1-713) | Fig S9 |
| Trigger | pcDNA3.1-SpCas9(714-1368) | https://benchling.com/s/seq-EgutSzE4sOjkDFpvAga6 | SpCas9(714-1368) | Fig S9 |
| Trigger | pcDNA3.1-SpCas9(1-713)-N_intein | https://benchling.com/s/seq-LznvtjWCo3gRuQTQLrFV | N-Cas9 | Fig 2 |
| Trigger | pcDNA3.1-C_intein-SpCas9(714-1368) | https://benchling.com/s/seq-IQZPP14tzFCrLOAcMehv | C-Cas9 | Fig 2 |
| Trigger | pcDNA3.1-SpCas9(1-713)-(GGGGS)3-DmrA | https://benchling.com/s/seq-dv1jMt9Ay11JeCJLUQVH | | Fig 2 |
| Trigger | pcDNA3.1-DmrC-(GGGGS)3-SpCas9(714-1368) | https://benchling.com/s/seq-Fm4hweFErSPt3TEBtD2O | | Fig 2 |
| Acr | pcDNA3.1+-AcrIIC2_Nm | https://benchling.com/s/seq-q7VMJRLxbI80KzAUR0GI | AcrIIC2 | Fig 2 |

| | | | | |
|---|---|---|---|---|
| OFF switch | pGluc-Sa_gRNA-tagRFP | https://benchling.com/s/seq-CdGyAKcK4BLcaxnwoJqB | | Fig S13 |
| TX-TL | pGluc-Sp_gRNA-tagRFP | https://benchling.com/s/seq-lDi3fB9tIGStqMMVnkq5 | pSp_gRNA-RFP or Sp_gRNA-RFP | Fig 4 |
| TX-TL | pTRE-Tight-hmAG1 | https://benchling.com/s/seq-2cqCEKIUhTzatrtUhv0P | pTRE-hmAG1 | Fig 4 |
| TX-TL | SP-dCas9-VPR | https://benchling.com/s/seq-ZEIwRvTTCW7jt0rU9sK6 | dSpCas9-VPR | Fig 4, 6 |
| TX-TL | pHL-gRNA[TRE new]-iRFP-RIH | https://benchling.com/s/seq-q7JGwy2sJVyYMaulUoN3 | gRNA | Fig 4 |
| 60 AND | pAkCas12b_sgRNA_v1-hMbCpf1 | https://benchling.com/s/seq-nCZ2RO20R6AumIh4YjgK | | Fig5 |
| 60 AND | pAkCas12b_sgRNA_v1-NcCas9 | https://benchling.com/s/seq-1a7DesFIHNILsHpSu4w9 | | Fig5 |
| 60 AND | pAkCas12b_sgRNA_v1-PguCas13b-NES | https://benchling.com/s/seq-B67ojjSjELQKJys2zkGN | | Fig5 |
| 60 AND | pAkCas12b_sgRNA_v1-PspCas13b-NES | https://benchling.com/s/seq-qz4qIieBfHwUzOWjC5e0 | | Fig5 |
| 60 AND | pAkCas12b_sgRNA_v1-SaCas9 | https://benchling.com/s/seq-bzPeWMvq81ItvXucSkKQ | | Fig5 |
| 60 AND | pGluc-Pgu_crRNA-AkCas12b | https://benchling.com/s/seq-9jNJRU5L8trZNL9Ea1Fk | | Fig5 |
| 60 AND | pGluc-Pgu_crRNA-hMbCpf1 | https://benchling.com/s/seq-K4EDhN81LGr1ZKzR7Zv4 | | Fig5 |
| 60 AND | pGluc-Pgu_crRNA-NcCas9 | https://benchling.com/s/seq-aLCsxbyWI32xJbcDvUvE | | Fig5 |
| 60 AND | pGluc-Pgu_crRNA-PspCas13b | https://benchling.com/s/seq-SjvgeWN3ybsNDiRJL3bI | | Fig5 |
| 60 AND | pGluc-Pgu_crRNA-SaCas9 | https://benchling.com/s/seq-xwARaxvsdDQ0dPAR5yLI | | Fig5 |
| 60 AND | pGluc-Psp_crRNA-AkCas12b | https://benchling.com/s/seq-6uVhkXlqnt3fl0fTPK9b | | Fig5 |
| 60 AND | pGluc-Psp_crRNA-hMbCpf1 | https://benchling.com/s/seq-exsrOWx6k6to67kEyp9l | | Fig5 |
| 60 AND | pGluc-Psp_crRNA-NcCas9 | https://benchling.com/s/seq-RfV6xht046zxIi5hD6kJ | | Fig5 |
| 60 AND | pGluc-Psp_crRNA-PguCas13b-NES | https://benchling.com/s/seq-ErjJA7ofbgNchGonVhRw | | Fig5 |
| 60 AND | pGluc-Psp_crRNA-SaCas9 | https://benchling.com/s/seq-vMQG9zSxZXBfRrJbHaUu | | Fig5 |
| 60 AND | pGluc-Sa_gRNA-AkCas12b | https://benchling.com/s/seq-gsFRzqnDG9GMjHBUxcXK | | Fig5 |
| 60 AND | pGluc-Sa_gRNA-MbCas12a | https://benchling.com/s/seq-pcMlOudTctmDddOgJqYx | | Fig5 |
| 60 AND | pGluc-Sa_gRNA-NcCas9 | https://benchling.com/s/seq-PKfTqfWkEK0qDtiR6NaT | | Fig5 |
| 60 AND | pGluc-Sa_gRNA-PguCas13b-NES | https://benchling.com/s/seq-1nJKeWZjZPKnwLvLAXw7 | | Fig5 |
| 60 AND | pGluc-Sa_gRNA-PspCas13b-NES | https://benchling.com/s/seq-vZdLDuRUj7lEkoSsrUus | | Fig5 |
| 60 AND | pMbCas12a_crRNA-AkCas12b | https://benchling.com/s/seq-ij4dn2iIGnGvaZlxhdfJ | | Fig5 |
| 60 AND | pMbCas12a_crRNA-NcCas9 | https://benchling.com/s/seq-blf5FyEY59ZpzSoStpet | | Fig5 |
| 60 AND | pMbCas12a_crRNA-PguCas13b-NES | https://benchling.com/s/seq-3xdNZDrleFaTH8daggKG | | Fig5 |
| 60 AND | pMbCas12a_crRNA-PspCas13b-NES | https://benchling.com/s/seq-qHKPavIjaz79DIq4Mk5f | | Fig5 |

| | | | | |
|---|---|---|---|---|
| 60 AND | pMbCas12a_crRNA-SaCas9 | https://benchling.com/s/seq-n012CwEHH9uvfaQ0gvj5 | | Fig5 |
| 60 AND | pNc_gRNA_v1-AkCas12b | https://benchling.com/s/seq-uM9EVPlutz8XRqTR19bt | | Fig5 |
| 60 AND | pNc_gRNA_v1-MbCpf1 | https://benchling.com/s/seq-PY2fTqoxv4KsMFs6I2GF | | Fig5 |
| 60 AND | pNc_gRNA_v1-PguCas13b-NES | https://benchling.com/s/seq-HdyvKPFRGxGzxA18WgRv | | Fig5 |
| 60 AND | pNc_gRNA_v1-PspCas13b-NES | https://benchling.com/s/seq-QHZAHBtl1644ZiJJs246 | | Fig5 |
| 60 AND | pNc_gRNA_v1-SaCas9 | https://benchling.com/s/seq-uO9Q7cbMhicDBiKJg0l8 | | Fig5 |
| Half-subtractor | pSa_IgRNA_a-CMVmin-Gluc_Sp_gRNA-tagBFP-Triplex-HHR-Sa_gRNA[TRE]-HDVR | https://benchling.com/s/seq-uf65bSQ0kDL47n4kCUnW | | Fig6 |
| Half-subtractor | pSp_IgRNA_a-CMVmin-Gluc_Sa_gRNA-tagBFP | https://benchling.com/s/seq-alXjZwBUovuZrMfD2Xm3 | | Fig6 |
| Half-subtractor | pSp_IgRNA_ax2-CMVmin-Gluc_Sa_gRNA-tagBFP | https://benchling.com/s/seq-KpJNkbiOwhNvphA7kTqG | | Fig6 |
| Half-subtractor | pTRE-Tight-Gluc_Sp_gRNA-hmAG1 | https://benchling.com/s/seq-RtHU0UdngTZHkwAhHqPP | | Fig6 |
| Half-subtractor | pcDNA3.1+-dSaCas9-VPR | https://benchling.com/s/seq-VOVsrV43YHMX6DoRTtEC | | Fig6 |
| Half-subtractor | pHL-Sa_IgRNA_a-iRFP-RIH | https://benchling.com/s/seq-Ay5dnk1zINPfqphQkY9X | | Fig6 |
| Half-subtractor | pHL-Sp_IgRNA_a-iRFP-RIH | https://benchling.com/s/seq-BNfF7C046rnblBDNHpsC | | Fig6 |
| | pcDNA3.1+-myc-HisA | https://benchling.com/s/seq-wZkotHbe8PB30KDVPJqa | Control/ No trigger | |

**Supplementary Table 4**

Primers, template oligo DNA for generating synthetic mRNAs.

PCR primers

| No. | Primer/Oligo DNA name | Sequence (5' → 3') |
|---|---|---|
| 1 | TAP_T7_G3C fwd primer | CAGTGAATTGTAATACGACTCACTATAGGGC |
| 2 | IVT_5prime_UTR primer | CAGTGAATTGTAATACGACTCACTATAGGGCGAATTAAGAGAGAAAAGAAGAGTAAGAAGAAATATA AGACACCGGTCGCCACCATG |
| 3 | Rev5UTR primer | CATGGTGGCGACCGGTGTCTTATATTTCTTCTTACTC |
| 4 | IVT_3prime_UTR primer | TCTAGACCTTCTGCGGGGCTTGCCTTCTGGCCATGCCCTTCTTCTCTCCCTTGCACCTGTACCTCTT GGTCTTTGAATAAAGCCTGAGTAGG |
| 5 | Fwd3UTR primer | TCTAGACCTTCTGCGGGGC |
| 6 | Rev3UTR2T20 | TTTTTTTTTTTTTTTTTTTTCCTACTCAGGCTTTATTCAAAGACCAAG |
| 7 | SphcCas9 ORF fwd primer | CACCGGTCGCCACCATGGATAAGAAATACAGCATTGGAC |
| 8 | SphcCas9 ORF rev primer | GCCCCGCAGAAGGTCTAGACTATCACACCTTCCTCTTCTTCTTGG |
| 9 | T7-5UTR for cassette1 | TAATACGACTCACTATAggTCAGATCCGCTAGCGGATCC |
| 10 | EGFP ORF_Rv | GCCCCGCAGAAGGTCTAGACTACTTGTACAGCTCGTCCATGCCGAGAG |
| 11 | EGFP mRNA1 ORF_Fw | CACCGGTCGCCACCATGGTGAGCAAGGGCGAG |
| 12 | EGFP mRNA1 ORF_Rv | CCTACTCAGGCTTTATTCATGTGAAATTTGTGATGCTATTGCTTTATTTGTAAC |
| 13 | YF771_T7_5UTR_fwd | ATTGTAATACGACTCACTATAGGGCGAATTAAGAGAGAAAAGAAGAGTAAG |
| 14 | 3UTR120A | TTTTTTTTTTTTTTTTTTTTTTTTTTTTTTTTTTTTTTTTTTTTTTTTTTTTTTTTTTTTTTTTTTTTTTTTTTTT TTTTTTTTTTTTTTTTTTTTTTTTTTTTTTTTTTTTTTTTTTTTCCTACTCAGGCTTTATTCA |

Key PCR products

| No. | Name | Type | Templates/Plasmids | Forward primer | Reverse primer |
|---|---|---|---|---|---|
| 1 | 5'-UTR | UTR | IVT_5prime_UTR primer | TAP_T7_G3C fwd primer | Rev5UTR primer |
| 2 | 3'-UTR | UTR | IVT_3prime_UTR primer | Fwd3UTR primer | Rev3UTR2T20 |
| 3 | Cas9 mRNA ORF | ORF | pHL-EF1a-SphcCas9-iC-A | SphcCas9 ORF fwd primer | SphcCas9 ORF rev primer |
| 5 | Gluc-Sp_gRNA-EGFP ORF | 5'UTR-ORF | pGluc-Sp_gRNA-EGFP | T7-5UTR for cassette1 | EGFP ORF_Rv |
| 7 | EGFP mRNA ORF | 5'UTR-ORF | pAptamerCassette-EGFP | EGFP mRNA1 ORF_Fw | EGFP mRNA1 ORF_Rv |
| 4 | Cas9 mRNA Template | IVT template | Cas9 mRNA ORF, 5'-UTR, 3'-UTR | TAP_T7_G3C fwd primer | 3UTR120A |
| 6 | Gluc-Sp_gRNA-EGFP Template | IVT template | Gluc-Sp_gRNA-EGFP ORF, 3'-UTR | T7-5UTR for cassette1 | 3UTR120A |
| 8 | EGFP mRNA Template | IVT template | EGFP mRNA ORF, 5'-UTR | TAP_T7_G3C fwd primer | 3UTR120A |
| 9 | iRFP670 mRNA Template | IVT template | pUC19-iRFP670woT7f | YF771_T7_5UTR_fwd | 3UTR120A |

**Supplementary Table 5**
Transfection tables of all experiments performed in this study.

Figure 1, Figure S2, S3A, S5B, S6, S7 (24-well plate)

| | |
|---|---|
| Switch plasmid | 100 ng |
| Trigger plasmid | 400 ng |
| Reference plasmid | 100 ng |
| Opti-MEM | up to 100 μL |
| Lipofectamine 2000 | 2 μL |

Figure 2B (24-well plate)

| | [N-Cas9, C-Cas9] [-, -] | [N-Cas9, C-Cas9] [+, -] | [N-Cas9, C-Cas9] [-, +] | [N-Cas9, C-Cas9] [+, +] | WT |
|---|---|---|---|---|---|
| pGluc-Sp_gRNA-EGFP | 100 ng | 100 ng | 100 ng | 100 ng | 100 ng |
| pcDNA3.1-myc-His6 | 800 ng | 400 ng | 400 ng | | 400 ng |
| pcDNA3.1-SpCas9 | | | | | 400 ng |
| pcDNA3.1-SpCas9(1-713)-N_intein | | 400 ng | | 400 ng | |
| pcDNA3.1-C_intein-SpCas9(714-1368) | | | 400 ng | 400 ng | |
| pCMV-tdiRFP670 | 100 ng | 100 ng | 100 ng | 100 ng | 100 ng |

Figure 2D (24-well plate)

| | A/C heterodimerizer (-) | | | | | |
|---|---|---|---|---|---|---|
| | WT | | Split | | No trigger | |
| | No_gRNA | Sp_gRNA | No_gRNA | Sp_gRNA | No_gRNA | Sp_gRNA |
| pAptamerCassette-EGFP | 100 ng | | 100 ng | | 100 ng | |
| pGluc-Sp_gRNA-EGFP | | 100 ng | | 100 ng | | 100 ng |
| pcDNA3.1-SpCas9(1-713)-(GGGGS)3-DmrA | | | 400 ng | 400 ng | | |
| pcDNA3.1-DmrC-(GGGGS)3-SpCas9(714-1368) | | | 400 ng | 400 ng | | |
| pcDNA3.1-myc-His6 | 400 ng | 400 ng | | | 800 ng | 800 ng |
| pcDNA3.1-SpCas9 | 400 ng | 400 ng | | | | |
| | A/C heterodimerizer (+) | | | | | |
| | WT | | Split | | No trigger | |
| | No_gRNA | Sp_gRNA | No_gRNA | Sp_gRNA | No_gRNA | Sp_gRNA |
| pAptamerCassette-EGFP | 100 ng | | 100 ng | | 100 ng | |
| pGluc-Sp_gRNA-EGFP | | 100 ng | | 100 ng | | 100 ng |
| pcDNA3.1-SpCas9(1-713)-(GGGGS)3-DmrA | | | 400 ng | 400 ng | | |
| pcDNA3.1-DmrC-(GGGGS)3-SpCas9(714-1368) | | | 400 ng | 400 ng | | |
| pcDNA3.1-myc-His6 | 400 ng | 400 ng | | | 800 ng | 800 ng |
| pcDNA3.1-SpCas9 | 400 ng | 400 ng | | | | |

Figure 2F (24-well plate)

| | 0 ng | 200 ng | 400 ng | 1000 ng | 2000 ng |
|---|---|---|---|---|---|
| pGluc-Sa_gRNA-EGFP or pAptamerCassette-EGFP | 100 ng | 100 ng | 100 ng | 100 ng | 100 ng |
| pcDNA3.1-SaCas9 | 200 ng | 200 ng | 200 ng | 200 ng | 200 ng |
| pcDNA3.1-myc-His6 | 2000 ng | 1000 ng | 400 ng | 200 ng | 0 ng |
| pcDNA3.1+-AcrIIC2_Nm | 0 ng | 200 ng | 400 ng | 1000 ng | 2000 ng |
| pCMV-tdiRFP670 | 100 ng | 100 ng | 100 ng | 100 ng | 100 ng |

Figure 3, S10-S12 (384-well plate)

| | |
|---|---|
| Switch plasmid | 31.25 ng |
| Trigger plasmid | 125 ng |
| Reference plasmid | 31.25 ng |
| Opti-MEM | up to 10 $\mu$L |
| Lipofectamine 2000 | 0.4 $\mu$L |

Figure 4 (24-well plate)

| notation | plasmid name | |
|---|---|---|
| dSpCas9-VPR | Sp_dCas9-VPR or pcDNA3.1+-myc-HisA | 400 [ng] |
| gRNA | pHL-gRNA[TRE new]-iRFP-RIH or pHL-Sp_IgRNA_Nluc-iRFP-RIH | 100 [ng] |
| pTRE-hmAG1 | pTRE-Tight-hmAG1 | 100 [ng] |
| pSp_gRNA-RFP | pGluc-Sp_gRNA-tagRFP or pcDNA3.1+-myc-HisA | 100 [ng] |
| reference | pAptamerCassette-tagBFP | 100 [ng] |

Figure 5 (96-well plate)

| | |
|---|---|
| Trigger plasmid 1 | 200 ng |
| Trigger plasmid 2 | 200 ng |
| Mediator plasmid 1 | 50 ng |
| Mediator plasmid 2 | 50 ng |
| Reporter plasmid | 12.5 ng |
| Reference plasmid | 25 ng |
| Opti-MEM | up to 20 $\mu$L |
| Lipofectamine 2000 | 0.5 $\mu$L |

| Figure 6 (24-well plate) | [0, 0] | [0, 1] | [1, 0] | [1, 1] |
|---|---|---|---|---|
| pTRE-Gluc_Sa_gRNA-hmA1 | 400 ng | | | |
| pHL-Sp_lgRNA_a-iRFP-RIH | 100 ng | | | |
| pHL-Sa_lgRNA_a-iRFP-RIH | 100 ng | | | |
| pcDNA3.1-myc-HisA | 800 ng | 400 ng | 400 ng | 0 ng |
| pcDNA3.1-dSpCas9-VPR | 0 ng | 400 ng | 0 ng | 400 ng |
| pcDNA3.1-dSaCas9-VPR | 0 ng | 0 ng | 400 ng | 400 ng |
| pSa_lgRNA_a-CMVmin-Gluc_Sp_gRNA-tagBFP | 100 ng | | | |
| pSp_lgRNA_ax2-CMVmin-Gluc_Sa_gRNA-tagBFP-Tri-HHR-Sp_gRNA[TRE]-HDVR | 400 ng | | | |
| Opti-MEM | Up to 100 $\mu$L | | | |
| Lipofectamine 2000 | 2 $\mu$L | | | |

Figure S3C (24-well plate)

| | |
|---|---|
| Switch/control plasmid | 100 ng |
| Trigger plasmid | 100 ng |
| Acr (AcrIIA4) plasmid | 100 ng |
| Reference plasmid | 100 ng |
| Opti-MEM | up to 20 $\mu$L |
| Lipofectamine 2000 | 0.5 $\mu$L |

Figure S4 (24-well plate)

| | |
|---|---|
| Switch/control mRNA | 100 ng |
| Trigger mRNA | 100 ng |
| Reference mRNA | 100 ng |
| Opti-MEM | up to 50 $\mu$L |
| Lipofectamine MessengerMax | 1 $\mu$L |

Figure S9 (24-well plate)

| | [N-Cas9, C-Cas9] [-, -] | [N-Cas9, C-Cas9] [+, -] | [N-Cas9, C-Cas9] [-, +] | [N-Cas9, C-Cas9] [+, +] | WT |
|---|---|---|---|---|---|
| pGluc-Sp_gRNA-EGFP | 100 ng | 100 ng | 100 ng | 100 ng | 100 ng |
| pcDNA3.1-myc-His6 | 800 ng | 400 ng | 400 ng | | 400 ng |
| pcDNA3.1-SpCas9 | | | | | 400 ng |
| pcDNA3.1-SpCas9(1-713) | | 400 ng | | 400 ng | |
| pcDNA3.1-SpCas9(714-1368) | | | 400 ng | 400 ng | |
| pCMV-tdiRFP670 | 100 ng | 100 ng | 100 ng | 100 ng | 100 ng |

Figure S13 (24-well plate)

| notation | plasmid name | |
|---|---|---|
| OFF switch | pGluc-Sa_gRNA-tagRFP<br>or<br>pAptamerCassette-tagRFP2 | 100 ng |
| ON switch | pNMD-ON-Gluc-Sa_gRNA<br>or<br>pNMD-ON-Gluc-Sa_gRNA | 100 ng |
| trigger | pcDNA3.1＋-SaCas9<br>or<br>pcDNA3.1＋-myc-HisA | 400 ng |
| reference | pCMV-tdiRFP | 100 ng |

**Supplementary Sequences**

RNA sequences used in this study.

1. The 5’ terminus of mRNA is capped with ARCA.
2. The protein coding regions are shown as bold letters.
3. The start sites and stop codons are underlined.

Cas9 mRNA

GGGCGAAUUAAGAGAGAAAAGAAGAGUAAGAAGAAAUAUAAGACACCGGUCGCCACC**AUGGAUAAGAAAUAC**
**AGCAUUGGACUGGACAUUGGGACAAACUCCGUGGGAUGGGCCGUGAUUACAGACGAAUACAAAGUGCCUU**
**CAAAGAAGUUCAAGGUGCUGGGCAACACCGAUAGACACAGCAUCAAGAAAAAUCUGAUUGGAGCCCUGCUG**
**UUCGACUCCGGCGAGACAGCUGAAGCAACUCGGCUGAAAAGAACUGCUCGGAGAAGGUAUACCCGCCGAA**
**AGAAUAGGAUCUGCUACCUGCAGGAGAUUUUCAGCAACGAAAUGGCCAAGGUGGACGAUAGUUUCUUUCAC**
**CGCCUGGAGGAAUCAUUCCUGGUCGAGGAAGAUAAGAAACACGAGCGGCAUCCCAUCUUUGGCAACAUUG**
**UGGACGAGGUCGCUUAUCACGAAAAGUACCCUACCAUCUAUCAUCUGAGGAAGAAACUGGUGGACUCCACA**
**GAUAAAGCAGACCUGCGCCUGAUCUAUCUGGCCCUGGCUCACAUGAUUAAGUUCCGGGGCCAUUUUCUGAU**
**CGAGGGGGAUCUGAACCCAGACAAUUCUGAUGUGGACAAGCUGUUCAUCCAGCUGGUCCAGACAUACAAUC**
**AGCUGUUUGAGGAAAACCCCAUUAAUGCAUCUGGCGUGGACGCAAAAGCCAUCCUGAGUGCCAGACUGUCU**
**AAGAGUCGGAGACUGGAGAACCUGAUCGCUCAGCUGCCAGGGGAAAAGAAAAACGGCCUGUUUGGGAAUC**
**UGAUUGCACUGUCACUGGGACUGACUCCCAACUUCAAGAGCAAUUUUGAUCUGGCCGAGGACGCUAAACUG**
**CAGCUGUCCAAGGACACCUAUGACGAUGACCUGGAUAACCUGCUGGCUCAGAUCGGGGAUCAGUACGCAG**
**ACCUGUUCCUGGCCGCUAAGAAUCUGUCUGACGCCAUCCUGCUGAGUGAUAUUCUGCGCGUGAACACCGAG**
**AUUACAAAAGCCCCCCUGUCAGCUAGCAUGAUCAAGAGAUAUGACGAGCACCAUCAGGAUCUGACCCUGCU**
**GAAGGCUCUGGUGAGGCAGCAGCUGCCUGAGAAGUACAAGGAAAUCUUCUUUGAUCAGUCUAAGAACGGA**
**UACGCCGGCUAUAUUGACGGCGGGGCUAGUCAGGAGGAGUUCUACAAGUUUAUCAAACCCAUUCUGGAGA**
**AGAUGGAUGGCACAGAGGAACUGCUGGUGAAACUGAAUCGGGAAGACCUGCUGAGGAAGCAGCGCACUUU**
**UGAUAACGGAAGCAUCCCUCACCAGAUUCAUCUGGGAGAGCUGCACGCAAUCCUGAGGCGCCAGGAAGAC**
**UUCUACCCAUUUCUGAAGGAUAACAGGGAGAAGAUCGAAAAAAUUCUGACAUUCCGCAUCCCCUACUAUGU**
**GGGCCCUCUGGCAAGAGGCAACAGCCGGUUUGCCUGGAUGACUCGCAAAUCUGAGGAAACAAUCACUCCC**
**UGGAACUUCGAGGAAGUGGUCGAUAAGGGCGCUUCCGCACAGUCUUUCAUUGAGCGGAUGACAAACUUCG**
**ACAAGAACCUGCCAAACGAAAAAGUGCUGCCCAAGCACUCUCUGCUGUACGAGUAUUUCACAGUCUAUAAC**
**GAACUGACUAAGGUGAAAUACGUCACCGAGGGGAUGAGAAAGCCUGCCUUCCUGAGUGGAGAACAGAAGA**
**AAGCUAUCGUGGACCUGCUGUUUAAAACCAAUAGGAAGGUGACAGUCAAGCAGCUGAAAGAGGACUAUUUC**
**AAGAAAAUUGAAUGUUUCGAUUCUGUGGAGAUCAGUGGCGUCGAAGACAGGUUUAACGCCUCCCUGGGGA**
**CCUACCACGAUCUGCUGAAGAUCAUUAAGGAUAAAGACUUCCUGGACAACGAGGAAAAUGAGGAUAUCCUG**
**GAAGACAUUGUGCUGACCCUGACACUGUUUGAGGAUAGGGAAAUGAUCGAGGAACGCCUGAAGACCUAUG**
**CCCAUCUGUUCGAUGACAAAGUGAUGAAACAGCUGAAGCGACGGAGAUACACAGGAUGGGGCCGACUGUC**
**UCGGAAGCUGAUCAAUGGGAUUCGCGACAAACAGAGUGGAAAGACCAUCCUGGACUUUCUGAAAUCAGAU**
**GGCUUCGCCAACCGGAACUUCAUGCAGCUGAUUCACGAUGACAGCCUGACAUUCAAAGAGGAUAUCCAGAA**
**GGCACAGGUGUCCGGGCAGGGAGACUCUCUGCACGAGCAUAUCGCAAACCUGGCCGGCAGCCCUGCCAUC**
**AAGAAAGGGAUUCUGCAGACCGUGAAGGUGGUGGACGAGCUGGUGAAAGUCAUGGGAAGACAUAAGCCAG**
**AAAACAUCGUGAUUGAGAUGGCCAGGGAAAAUCAGACCACACAGAAAGGCCAGAAGAACUCAAGGGAGCG**
**CAUGAAAAGAAUCGAGGAAGGAAUUAAGGAACUGGGCAGCCAGAUCCUGAAAGAGCACCCCGUGGAAAAC**
**ACACAGCUGCAGAAUGAGAAGCUGUAUCUGUACUAUCUGCAGAAUGGACGCGAUAUGUACGUGGACCAGG**
**AGCUGGAUAUUAACCGACUGUCCGAUUACGACGUGGAUCAUAUCGUCCCACAGUCAUUCCUGAAAGAUGAC**
**AGCAUUGACAAUAAGGUGCUGACCCGCUCUGACAAAAACCGAGGCAAGAGUGAUAAUGUCCCCUCAGAGGA**
**AGUGGUCAAGAAAAUGAAGAACUACUGGAGGCAGCUGCUGAAUGCCAAACUGAUCACACAGCGAAAGUUU**
**GAUAACCUGACUAAAGCUGAGCGGGGAGGCCUGAGUGAACUGGACAAAGCAGGCUUCAUUAAGCGACAGC**
**UGGUGGAGACACGGCAGAUCACAAAGCACGUCGCCCAGAUUCUGGAUUCAAGAAUGAACACUAAGUACGAU**
**GAGAAUGACAAACUGAUCAGAGAAGUGAAGGUCAUUACCCUGAAGUCAAAACUGGUGAGCGACUUUCGGA**
**AAGAUUUCCAGUUUUAUAAGGUCAGAGAGAUCAACAACUACCACCAUGCUCAUGACGCAUACCUGAACGCA**
**GUGGUCGGCACAGCCCUGAUUAAGAAAUACCCUAAACUGGAGUCCGAGUUCGUGUACGGGGACUAUAAGG**
**UGUACGAUGUCAGAAAAAUGAUCGCCAAGUCUGAGCAGGAAAUUGGCAAAGCCACUGCUAAGUAUUUCUUU**
**UACAGUAACAUCAUGAAUUUCUUUAAGACUGAGAUCACCCUGGCAAAUGGGGAAAUCCGAAAGCGGCCACU**
**GAUUGAGACUAACGGCGAGACAGGAGAAAUCGUGUGGGACAAAGGAAGAGAUUUUGCUACCGUGAGGAAG**
**GUCCUGAGCAUGCCCCAAGUGAAUAUUGUCAAGAAAACAGAGGUGCAGACUGGGGGAUUCAGUAAGGAAU**
**CAAUUCUGCCUAAACGCAACUCCGAUAAGCUGAUCGCCCGAAAGAAAGACUGGGACCCCAAGAAGUAUGGC**
**GGGUUCGACUCCCCAACUGUGGCUUACUCUGUCCUGGUGGUCGCAAAGGUGGAGAAGGGAAAAAGCAAGA**
**AACUGAAAUCCGUCAAGGAACUGCUGGGCAUCACCAUUAUGGAGCGCAGCUCCUUCGAAAAGAAUCCUAUC**
**GAUUUUCUGGAGGCCAAAGGCUAUAAGGAAGUGAAGAAAGACCUGAUCAUCAAGCUGCCAAAGUACUCACU**
**GUUUGAGCUGGAAAACGGGAGAAAGAGGAUGCUGGCAAGCGCCGGGGAGCUGCAGAAAGGAAAUGAACUG**
**GCCCUGCCCUCCAAGUACGUGAACUUCCUGUAUCUGGCUAGCCACUACGAGAAGCUGAAAGGGUCCCCUGA**
**GGAUAACGAACAGAAACAGCUGUUUGUGGAGCAGCACAAGCAUUAUCUGGACGAGAUCAUUGAACAGAUUA**
**GCGAGUUCUCCAAAAGAGUGAUCCUGGCUGACGCAAAUCUGGAUAAGGUCCUGAGCGCAUACAACAAACAC**

**CGGGAUAAGCCAAUCAGAGAGCAGGCCGAAAAUAUCAUUCAUCUGUUCACUCUGACCAACCUGGGAGCCCC**
**CGCAGCCUUCAAGUAUUUGACACUACCAUCGAUCGCAAACGAUACACAAGCACUAAGGAGGUGCUGGACG**
**CUACCCUGAUUCAUCAGAGCAUUACUGGCCUGUAUGAAACAAGGAUUGACCUGUCUCAGCUGGGCGGCGAC**
**UCCGGAGCUGACCCCAAGAAGAAGAGGAAGGUGUGA**UAGUCUAGACCUUCUGCGGGGCUUGCCUUCUGGC
CAUGCCCUUCUUCUCUCCCUUGCACCUGUACCUCUUGGUCUUUGAAUAAAGCCUGAGUAGGAAAAAAAAAAAA
AAAAAAAAAAAAAAAAAAAAAAAAAAAAAAAAAAAAAAAAAAAAAAAAAAAAAAAAAAAAAAAAAAAAAAAAAAAAAAAAAAAAAAA
AAAAAAAAAAAAAAAAAAAAAAAAAAAAAAAAA

Gluc-Sp_gRNA-EGFP
GGUCAGAUCCGCUAGCGGAUCC*GAGAUCAGGGCAAACAGAACUGUUUUAGAGCUAGAAAUAGCAAGUUAAAA*
*UAAGGCUAGUCCGUUAUCAACUUGAAAAAGUGGCACCGAGUCGGUGC*AGAUCUACCGGUCGCCACC**AUGGUG**
**AGCAAGGGCGAGGAGCUGUUCACCGGGGUGGUGCCCAUCCUGGUCGAGCUGGACGGCGACGUAAACGGCC**
**ACAAGUUCAGCGUGUCCGGCGAGGGCGAGGGCGAUGCCACCUACGGCAAGCUGACCCUGAAGUUCAUCUG**
**CACCACCGGCAAGCUGCCCGUGCCCUGGCCCACCCUCGUGACCACCCUGACCUACGGCGUGCAGUGCUUCA**
**GCCGCUACCCCGACCACAUGAAGCAGCACGACUUCUUCAAGUCCGCCAUGCCCGAAGGCUACGUCCAGGAG**
**CGCACCAUCUUCUUCAAGGACGACGGCAACUACAAGACCCGCGCCGAGGUGAAGUUCGAGGGCGACACCC**
**UGGUGAACCGCAUCGAGCUGAAGGGCAUCGACUUCAAGGAGGACGGCAACAUCCUGGGGCACAAGCUGGA**
**GUACAACUACAACAGCCACAACGUCUAUAUCAUGGCCGACAAGCAGAAGAACGGCAUCAAGGUGAACUUCA**
**AGAUCCGCCACAACAUCGAGGACGGCAGCGUGCAGCUCGCCGACCACUACCAGCAGAACACCCCCAUCGGC**
**GACGGCCCCGUGCUGCUGCCCGACAACCACUACCUGAGCACCCAGUCCGCCCUGAGCAAAGACCCCAACGA**
**GAAGCGCGAUCACAUGGUCCUGCUGGAGUUCGUGACCGCCGCCGGGAUCACUCUCGGCAUGGACGAGCUG**
**UACAAG**UAGUCUAGACCUUCUGCGGGGCUUGCCUUCUGGCCAUGCCCUUCUUCUCUCCCUUGCACCUGUAC
CUCUUGGUCUUUGAAUAAAGCCUGAGUAGGAAAAAAAAAAAAAAAAAAAAAAAAAAAAAAAAAAAAAAAAAAAAAAAA
AAAAAAAAAAAAAAAAAAAAAAAAAAAAAAAAAAAAAAAAAAAAAAAAAAAAAAAAAAAAAAAAAAAAAAAAAAAAAAAAAAA

EGFP mRNA
GGCCGCUUGAAGUCUUUAAUUAAACCCGCUUGAAGUCUUUAAUUAAACGAACGGGCACGCUGACAAUUCAAG
CACUCUGAUUUGACAAUUACAAGCACUCUGAUUUGACAAUUACACCGGUCGCCACC**AUGGUGAGCAAGGGCG**
**AGGAGCUGUUCACCGGGGUGGUGCCCAUCCUGGUCGAGCUGGACGGCGACGUAAACGGCCACAAGUUCAG**
**CGUGUCCGGCGAGGGCGAGGGCGAUGCCACCUACGGCAAGCUGACCCUGAAGUUCAUCUGCACCACCGGC**
**AAGCUGCCCGUGCCCUGGCCCACCCUCGUGACCACCCUGACCUACGGCGUGCAGUGCUUCAGCCGCUACCC**
**CGACCACAUGAAGCAGCACGACUUCUUCAAGUCCGCCAUGCCCGAAGGCUACGUCCAGGAGCGCACCAUCU**
**UCUUCAAGGACGACGGCAACUACAAGACCCGCGCCGAGGUGAAGUUCGAGGGCGACACCCUGGUGAACCG**
**CAUCGAGCUGAAGGGCAUCGACUUCAAGGAGGACGGCAACAUCCUGGGGCACAAGCUGGAGUACAACUAC**
**AACAGCCACAACGUCUAUAUCAUGGCCGACAAGCAGAAGAACGGCAUCAAGGUGAACUUCAAGAUCCGCCA**
**CAACAUCGAGGACGGCAGCGUGCAGCUCGCCGACCACUACCAGCAGAACACCCCCAUCGGCGACGGCCCC**
**GUGCUGCUGCCCGACAACCACUACCUGAGCACCCAGUCCGCCCUGAGCAAAGACCCCAACGAGAAGCGCGA**
**UCACAUGGUCCUGCUGGAGUUCGUGACCGCCGCCGGGAUCACUCUCGGCAUGGACGAGCUGUACAAG**UAAA
GCGGCCGCGACUCUAGAUCAUAAUCAGCCAUACCACAUUUGUAGAGGUUUUACUUGCUUUAAAAAACCUCCCA
CACCUCCCCCUGAACCUGAAACAUAAAAUGAAUGCAAUUGUUGUUGUUAACUUGUUUAUUGCAGCUUAUAAUG
GUUACAAAUAAAGCAAUAGCAUCACAAAUUUCACAUGAAUAAAGCCUGAGUAGGAAAAAAAAAAAAAAAAAAAAA
AAAAAAAAAAAAAAAAAAAAAAAAAAAAAAAAAAAAAAAAAAAAAAAAAAAAAAAAAAAAAAAAAAAAAAAAAAAAAAAAAAAAAAA
AAAAAAAAAAAAAAAAAAAAAAA

iRFP670 mRNA
GGGCGAAUUAAGAGAGAAAAGAAGAGUAAGAAGAAAUAUAAGACACCGGUCGCCACC**AUGGCGCGUAAGGUC**
**GAUCUCACCUCCUGCGAUCGCGAGCCGAUCCACAUCCCCGGCAGCAUUCAGCCGUGCGGCUGCCUGCUAGC**
**CUGCGACGCGCAGGCGGUGCGGAUCACGCGCAUUACGGAAAAUGCCGGCGCGUUCUUUGGACGCGAAACU**
**CCGCGGGUCGGUGAGCUACUCGCCGAUUACUUCGGCGAGACCGAAGCCCAUGCGCUGCGCAACGCACUGG**
**CGCAGUCCUCCGAUCCAAAGCGACCGGCGCUGAUCUUCGGUUGGCGCGACGGCCUGACCGGCCGCACCUU**
**CGACAUCUCACUGCAUCGCCAUGACGGUACAUCGAUCAUCGAGUUCGAGCCUGCGGCGGCCGAACAGGCC**
**GACAAUCCGCUGCGGCUGACGCGGCAGAUCAUCGCGCGCACCAAAGAACUGAAGUCGCUCGAAGAGAUGG**
**CCGCACGGGUGCCGCGCUAUCUGCAGGCGAUGCUCGGCUAUCACCGCGUGAUGUUGUACCGCUUCGCGGA**
**CGACGGCUCCGGGAUGGUGAUCGGCGAGGCGAAGCGCAGCGACCUCGAGAGCUUUCUCGGUCAGCACUUU**
**CCGGCGUCGCUGGUCCCGCAGCAGGCGCGGCUACUGUACUUGAAGAACGCGAUCCGCGUGGUCUCGGAUU**
**CGCGCGGCAUCAGCAGCCGGAUCGUGCCCGAGCACGACGCCUCCGGCGCCGCGCUCGAUCUGUCGUUCGC**
**GCACCUGCGCAGCAUCUCGCCCUGCCAUCUCGAAUUUCUGCGGAACAUGGGCGUCAGCGCCUCGAUGUCG**
**CUGUCGAUCAUCAUUGACGGCACGCUAUGGGGAUUGAUCAUCUGUCAUCAUUACGAGCCGCGUGCCGUGC**
**CGAUGGCGCAGCGCGUCGCGGCCGAAAUGUUCGCCGACUUCUUAUCGCUGCACUUCACCGCCGCCCACCAC**
**CAACGCAGAUCUCAUAUGCAUCUCGAGUGA**UAGUCUAGACCUUCUGCGGGGCUUGCCUUCUGGCCAUGCCC
UUCUUCUCUCCCUUGCACCUGUACCUCUUGGUCUUUGAAUAAAGCCUGAGUAGGAAAAAAAAAAAAAAAAAAA
AAAAAAAAAAAAAAAAAAAAAAAAAAAAAAAAAAAAAAAAAAAAAAAAAAAAAAAAAAAAAAAAAAAAAAAAAAAAAAAAAAAAAAA
AAAAAAAAAAAAAAAAAAAAAAAAAA
